# Multiplexed code of navigation variables in anterior limbic areas

**DOI:** 10.1101/684464

**Authors:** Jean Laurens, Amada Abrego, Henry Cham, Briana Popeney, Yan Yu, Naama Rotem, Janna Aarse, Eftihia K. Asprodini, J. David Dickman, Dora E. Angelaki

## Abstract

The brain’s navigation system integrates multimodal cues to create a sense of position and orientation. Here we used a multimodal model to systematically assess how neurons in the anterior thalamic nuclei, retrosplenial cortex and anterior hippocampus of mice, as well as in the cingulum fiber bundle and the white matter regions surrounding the hippocampus, encode an array of navigational variables when animals forage in a circular arena. In addition to coding head direction, we found that some thalamic cells encode the animal’s allocentric position, similar to place cells. We also found that a large fraction of retrosplenial neurons, as well as some hippocampal neurons, encode the egocentric position of the arena’s boundary. We compared the multimodal model to traditional methods of head direction tuning and place field analysis, and found that the latter were inapplicable to multimodal regions such as the anterior thalamus and retrosplenial cortex. Our results draw a new picture of the signals carried and outputted by the anterior thalamus and retrosplenial cortex, offer new insights on navigational variables represented in the hippocampus and its vicinity, and emphasize the importance of using multimodal models to investigate neural coding throughout the navigation system.

## Introduction

One of the most striking properties of the rodent navigation system is the existence of characteristic cell populations that represent well-defined navigation variables. For instance, place cells are pyramidal neurons in the hippocampus (O’Keefe 1971, 1796) that encode the animal’s allocentric position, and head direction (HD) cells in the antero-dorsal thalamic nuclei form a neuronal compass (Taube et al. 1995; Blair and Sharp 1996; Zugaro et al. 2001; Peyrache et al. 2015, 2017; Page et al. 2017). Place cells and HD cells are thought the prominent neuron types in these regions, where no other navigational variables have been reported. In contrast, other regions such as the medial entorhinal cortex (MEC) encode multiple navigation variables and, as revealed by a recent study (Hardcastle et al. 2017), individual neurons often encode multiple variables. May this be a property exclusively for the MEC, or does mixed selectivity represent a generic property throughout the rodent spatial navigation circuit?

Here we used a Linear-Nonlinear (LN) model (Hardcastle et al. 2017) to characterize population responses in a network of brain regions involved in navigation. We first describe neurons in the anterior thalamic nuclei (ATN), and in particular the antero-dorsal thalamic nuclei, and whether they may encode spatially modulated variables other than head direction. We also test for the first time whether neuronal firing is phase-locked to theta-band LFP, a property that has never been reported for head direction cells (HDC) in the antero-dorsal thalamus, yet is frequent in antero-ventral thalamic nuclei HDC (Tsanov 2010, 2011).

Next, we compare neural selectivity in ATN and the retrosplenial cortex (RSC), noting few similarities. Unlike the ATN, we find that RSC is dominated by the coding of arena boundaries in an egocentric frame of reference (Alexander et al. 2019; Hinman et al. 2019; Gofman et al. 2019; see also Peyrache et al. 2017; Wang et al. 2018; Derdikman, 2009). We also characterized neuronal responses in anterior parts of the hippocampal formation (CA2/CA3). Furthermore, we also used the LN model to re-analyze previously published neuronal data recorded in the ATN (Peyrache et al. 2015). Finally, in order to test whether the multiplexed code is communicated among brain areas, we also describe response properties from the cingulum fiber bundle, in the vicinity of Bregma, i.e. at a location where it conveys output fibers from the ATN and RSC (Domesick 1970; van Groen and Vyss 1990, 1995, Bubb et al. 2017, 2018).

Traditional classification methods evaluate a cell’s response to a specific navigation variable by computing the corresponding tuning curve, without considering other variables. We found that these approaches produce biased results. In particular, head direction or egocentric boundary tuning is often mistaken for allocentric position tuning; to such an extent that attempting to detect place cells in the anterior thalamus or retrosplenial cortex by computing traditional “place fields” would result in excessive numbers of false positives. This stresses out the importance of using multiplexed models to systematically investigate brain regions where multiple variables are encoded.

## Results

We used tetrodes bundles to record extracellularly while mice (n=22) foraged in a circular arena (50 cm diameter; 8-minutes recording sessions) (**Fig. 1A**). We sampled 5 different brain regions (**Fig. 1B**): the anterior nuclei of the thalamus (ATN, n=7 animals), which included predominantly of recordings in the antero-dorsal nuclei; the retrosplenial cortex (RSC, n=4 animals); the cingulum fiber bundle (n=7 animals); the anterior portion of the hippocampus (CA2/CA3; n=3 animals); and the white matter region anterior to the hippocampus (Fimbria and fornix; n=6 animals). Multiple regions were sampled sequentially in some animals. Recording locations were verified post-mortem (**Fig. 1C**, see also **Fig. 6–10 S1**). We recorded and analyzed the responses of 1219 neurons (300 in ATN, 180 in RSC, 380 in the cingulum, 112 in hippocampus, and 247 in the fimbria).

**Figure 1:**
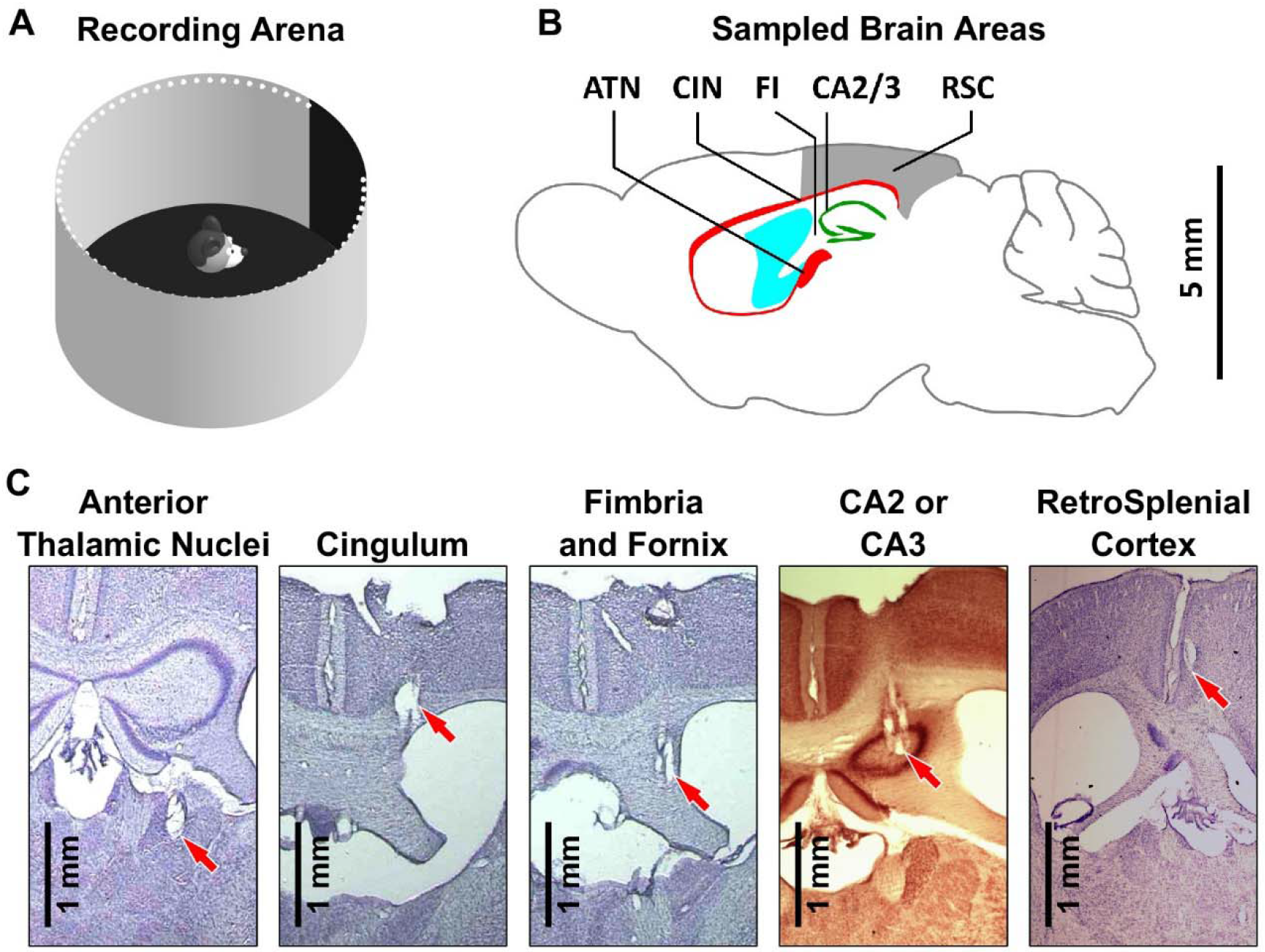
Overview of the experimental approach. **A:** Experimental apparatus. Mice move freely in a circular arena (50 cm in diameter). A black card provides an orienting cue. **B:** Sagittal section of a mouse brain at ^~^1 mm lateral of the midline, with all recorded regions indicated. ATN: Anterior Thalamic Nuclei. The cingulum fiber bundle (CIN) runs between the cortex and the corpus callosum. The fimbria and fornix (FI) refer here to white matter regions located around the anterior portion of the hippocampus. CA2/3: anterior portions of hippocampus. RSC: retrosplenial cortex. **C:** Representative coronal sections showing tetrode track marks (red arrows) in all recorded areas (see also **Fig. 6–10S1**).

We used a multivariate linear-nonlinear (LN) model (Hardcastle et al. 2017) to test whether each cell responds significantly to a series of navigational variables (**Fig. 2A**), and to assess whether it is modulated by theta-band (5-12 Hz) LFP. Specifically, we tested the 4 variables considered in (Hardcastle et al. 2017): the animal’s allocentric position (AP), head direction (HD) and linear speed (LS) in the environment, as well as the phase of the theta-band LFP rhythm (ΘP), and added two additional variables: the egocentric position of the arena’s boundary relative to the head (EB) and the head’s angular speed (AS). The LN model assumes that multiple variables influence the cell’s response in a multiplicative manner and uses an optimization procedure to compute each variable’s tuning curve to match the recorded firing rate, as shown next. To determine the best model, each cell’s response is fitted separately by each model variable alone, or with pairs of variables (or triplets, and so on) and uses a forward search procedure to determine which variables influence the cell’s firing significantly (according to the Log-likelihood ratio; see Methods).

**Figure 2:**
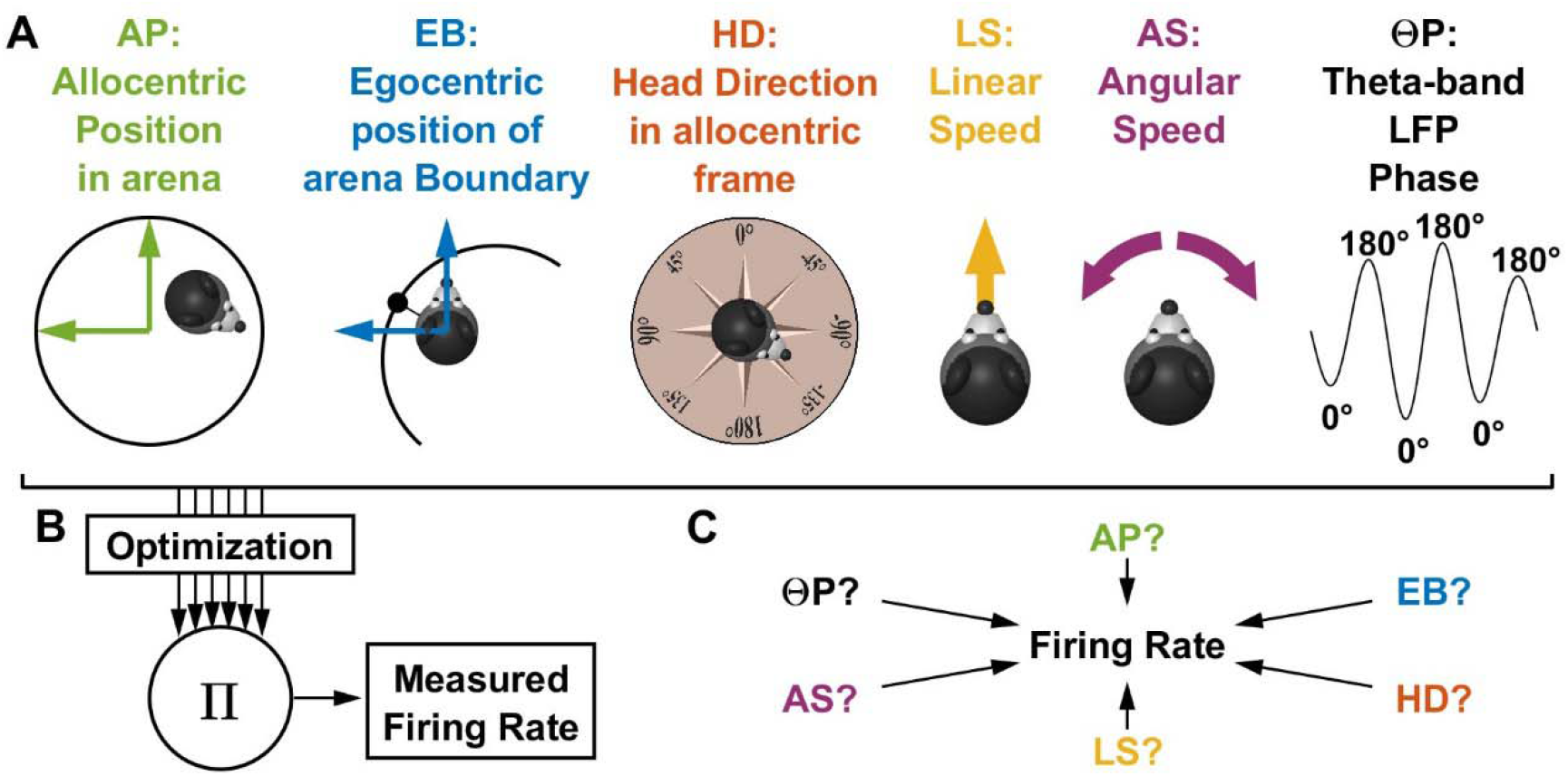
Overview of the LN model (Hardcastle et al. 2017). **A:** Representation of the 6 variables used in the LN model. The color code is used in subsequent figures. **B:** The model adjusts the tuning curve of all variables in order to fit the experimentally measured firing rate optimally, using a gradient ascent procedure, and assuming that the variables interact multiplicatively. **C:** A search procedure is used to determine which variables contribute significantly to the cell’s response (see Methods).

### Example cells

We first use two well-known neuron types to illustrate the robustness of the model: an example HD cell recorded in the ATN (**Fig. 3**); and an example hippocampal place cell (AP response) that also fired in phase with theta-LFP (ΘP modulation) (**Fig. 4**). Subsequently, we also illustrate examples of a novel cell type in RSC (**Fig. 5**), as well as coding for more multiplexed variables in other cell types.

**Figure 3:**
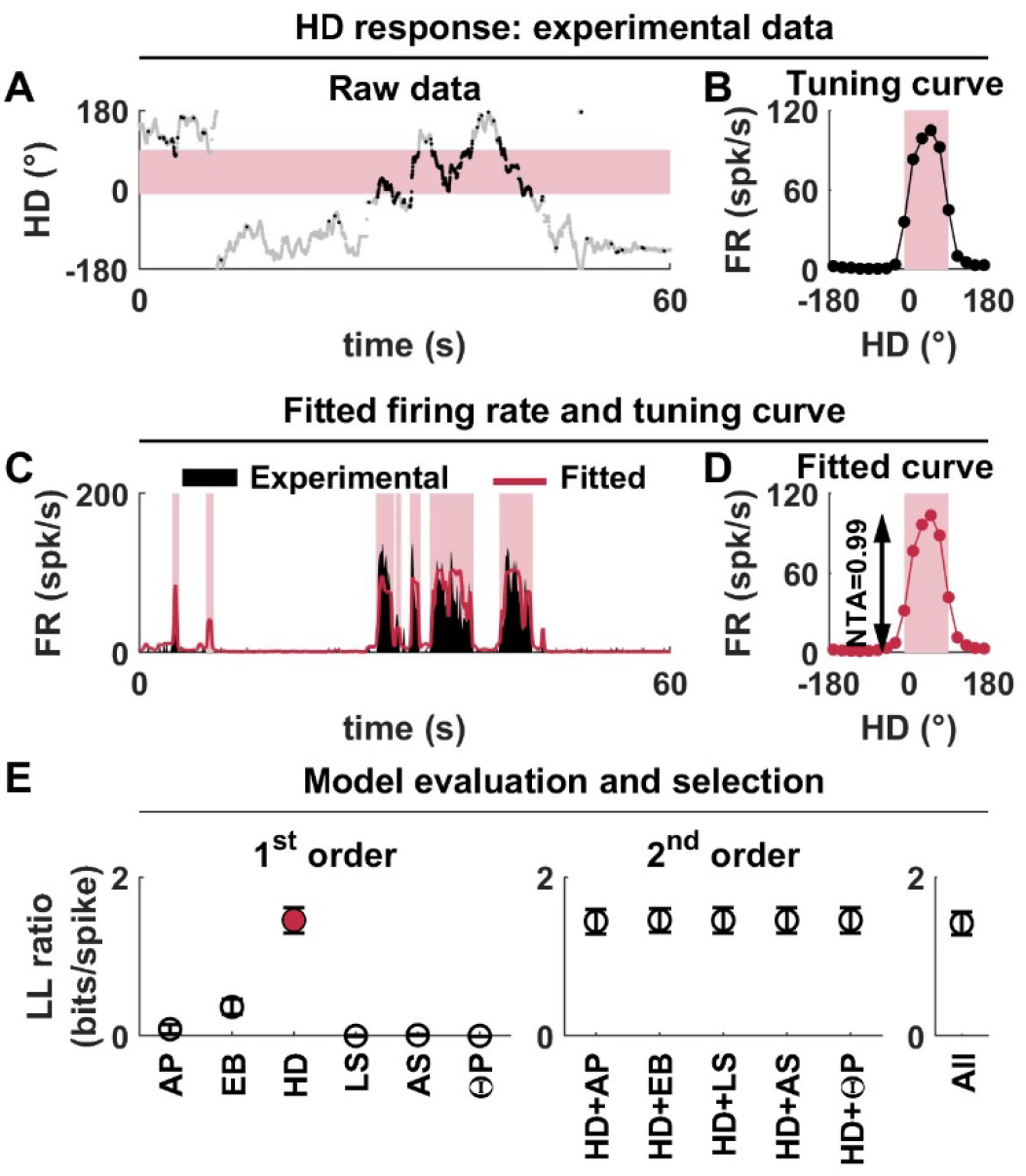
Example HD cell recorded in the ATN. **A:** Raw data. The orientation of the animal as a function of time is shown in grey, and recorded spikes are overlaid as black dots. The cell fires preferentially when the animal’s orientation is in the range of - 10° to 90°, which is indicated by a pink band. **B:** Traditional HD tuning curve computed by binning orientation data. The cell’s preferred firing range is indicated in pink. **C:** Model fit (red curve) to the cell’s firing rate (black). Time periods where HD is within the preferred firing range are indicated in red. The cell fires consistently during these time periods, and this firing is accurately reproduced by the model. **D:** HD tuning curve fitted by the LN model. In this simple example, the curve is identical to the experimental tuning curve in (B). Note, however, that this is not in general the case, as illustrated e.g. in Fig.5E. **E:** Illustration of the model selection procedure. Left panel: goodness of fit (LL ratio) of all 1^st^ order models. The goodness of fit of the HD model is higher than any other model. Middle panel: goodness of fit of all 2^nd^ order models. No model provides a significantly better fit than the HD model alone, which is therefore selected as the best model (red). Right panel: goodness of fit of the model including all variables, not significantly better than the HD model alone.

**Figure 4:**
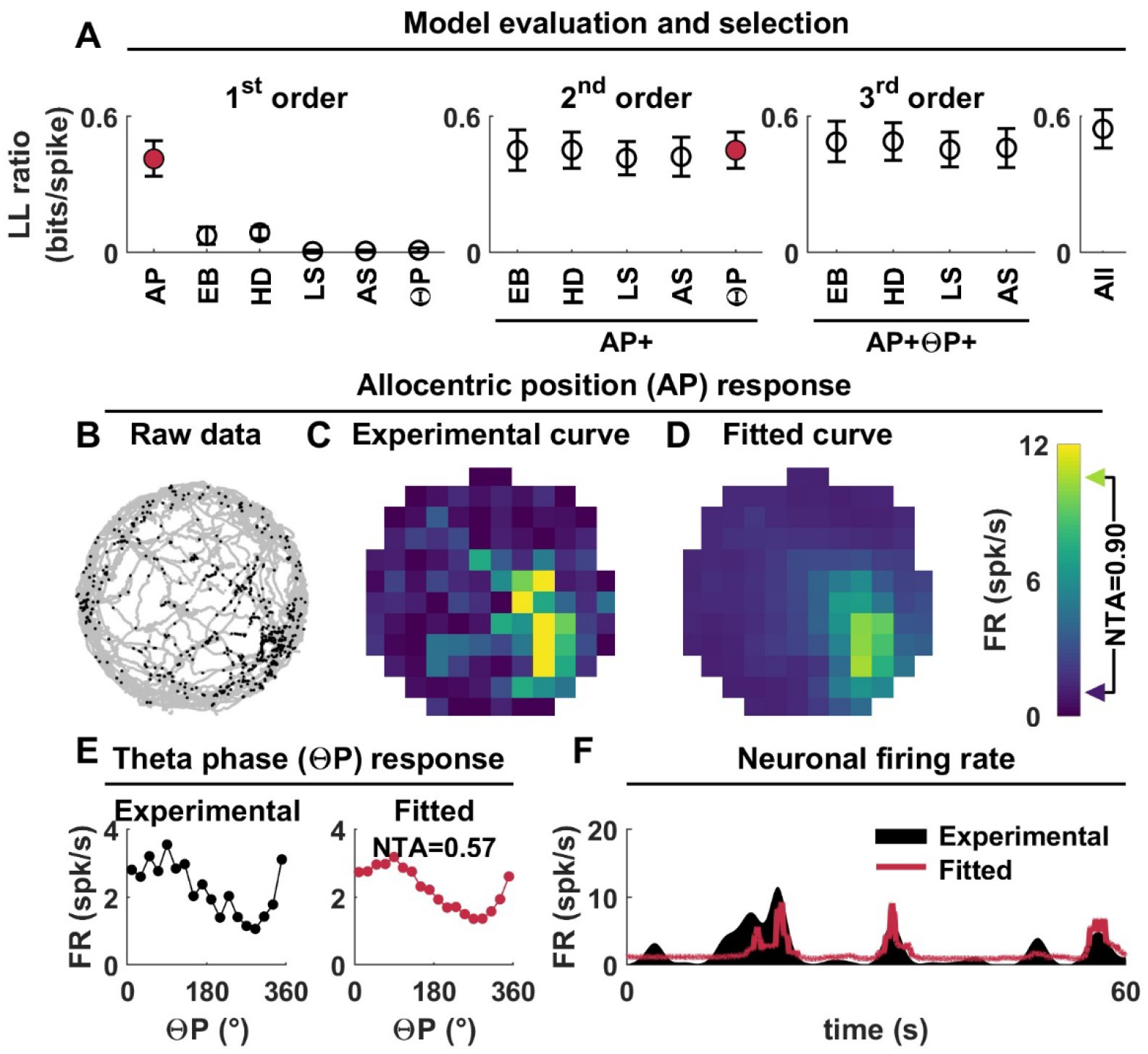
Example place cell recorded in the hippocampus. **A:** Illustration of the model selection procedure. The LN model tested a set of 1^st^, 2^nd^ and 3^rd^ order models and selected the AP+ΘP model (red). Therefore, this cell is modulated by the animal’s allocentric position (AP) and tends to fire in phase with theta-band LFP. **B:** Raw data. AP recorded over time is shown in grey, and recorded spikes are overlaid as black dots. **C:** Experimental AP tuning curve computed in the traditional way by binning the data and represented as a color map (the color scale is shown to the right of panel D). **D:** AP tuning curve fitted by the LN model. The curve resembles the experimental curve but is smoother due to the model’s smoothness constraint. **E:** Experimental (left) and fitted (right) ΘP tuning. The neuron responds preferentially at a phase of ^~^90°, i.e. between the peak and the trough of the LFP. **F:** Model fit (red curve) to a portion of the cell’s firing rate (black).

**Figure 5:**
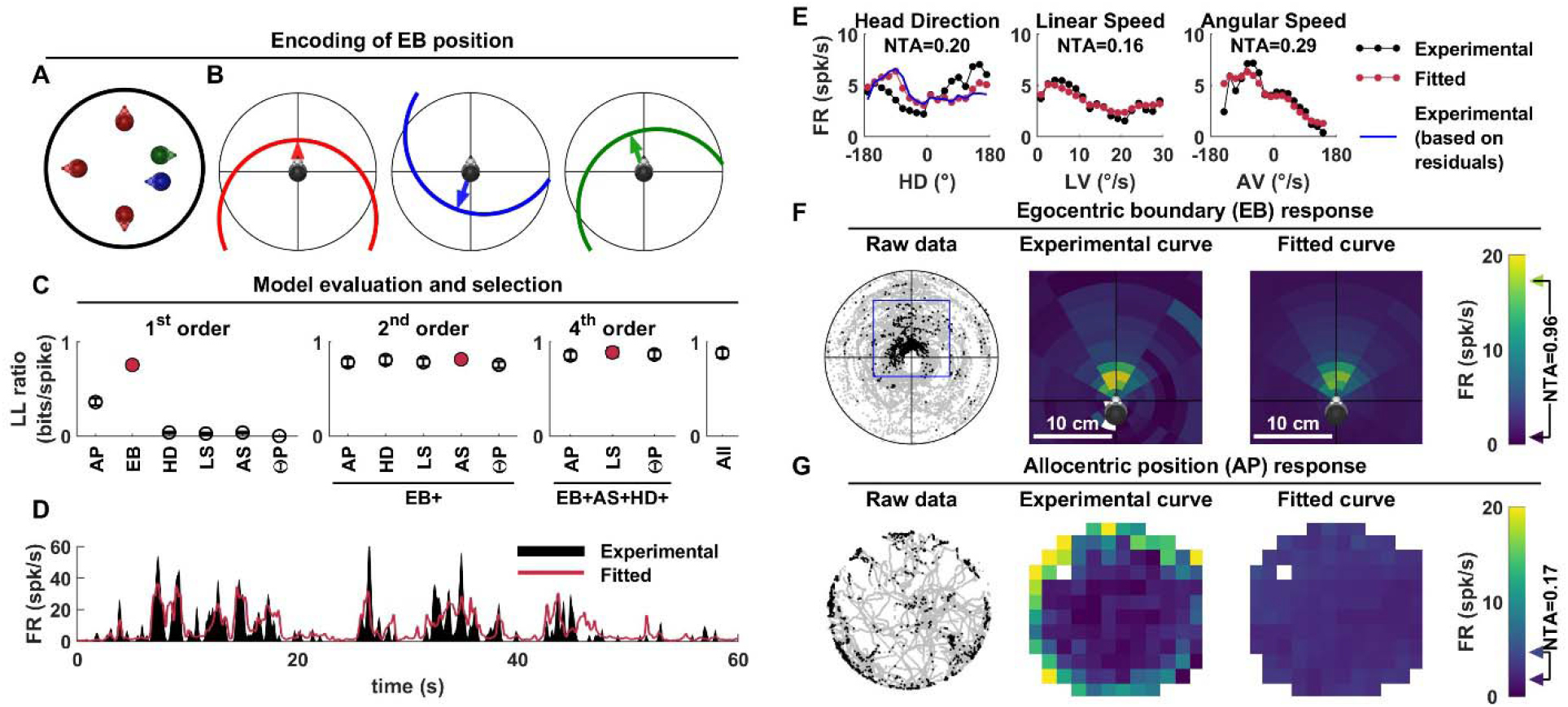
Example EBC recorded in RSC. **A:** Encoding of the egocentric position of the arena’s boundary. Five possible positions of the head within the arena are shown. The 3 heads colored in red are placed 10 cm away from the boundary and facing directly towards it. Therefore, from an egocentric perspective, their position relative to the arena are identical, although their allocentric positions are different. The green and blue colored heads are placed at a similar position, but with different orientations relative to the arena boundary. **B:** Egocentric positions corresponding to the allocentric positions for the example in A. Each panel corresponds to one example head position (or multiple equivalent positions for the first panel). The arena’s boundary is drawn relative to the head, and an arrow points towards its closet point. Because the arena is circular, and its radius is known, the position of the entire boundary can be known based on the position of the closest point of the boundary. Therefore, EB is represented as a 2D variable that encodes the position of the boundary closest point in egocentric coordinates (see **Suppl. Fig. 1** for details). **C:** Illustration of the model selection procedure. A set of models is tested until the 4^th^ order model EB+HD+AV+LS is selected as best fitting. **D:** Model fit (red curve) to the cell’s firing rate (black). The cell tends to fire in irregular bursts, some of which are not accurately predicted by the model, although the low-frequency response is consistently captured. **E:** Experimental (black) and fitted (red) HD (left), LS (middle) and AS (right) tuning curves. The experimental HD tuning curve (black) does not correspond to the fitted tuning curve (red) because the former is heavily biased by the cell’s sensitivity to EB. This can be demonstrated by fitting the EB model and re-computing the experimental HD tuning curve based on the residuals of that model. The resulting curve (blue) matches the fitted tuning curve closely. **F:** Example cell’s response to EB. Left: raw data, with EB encoded as shown in (B) and spikes overlaid as black dots (see also **Suppl. Movie** 1). Middle: experimental tuning curve. Right: fitted tuning curve. **G:** Example cell’s response to AP. Same legend as in Fig. 4A-C. Although the cell appears tuned to AP because it fires close to the boundarys, its response is in fact a result of its EB tuning. The fitted curve is flat, correctly reflecting the cell’s absence of significant AP response.

#### ATN example cell

The example HD cell fired in intense bursts when the head faced a specific allocentric direction (50°; **Fig. 3A,C**, black). We separated HD in 18 bins and computed the average firing rate within each bin. The resulting tuning curve (**Fig. 3B**) was typical of an ATN HD cell, exhibiting a sharp peak with an average firing of 104 spk/s at the preferred direction. Computing a HD tuning curve in this manner (the ‘experimental’ tuning curve) is the traditional approach for evaluating HD responses, typically combined with a shuffling-based statistical analysis of the curve’s tuning amplitude. By contrast, the LN model fits a tuning curve and uses a cross-validation procedure to test for statistical significance. In this cell, the ‘reconstructed’ tuning curve (**Fig. 3D**) resembles the experimental tuning curve (**Fig. 3C**).

To evaluate the set of variables encoded by this cell, we first fitted the firing rate based on each of the 6 variables individually (**Fig. 3E**, left). HD provided a considerably better fit, measured by the Log-likelihood ratio, than any other variable, and was selected as the best first-order model. Next, we tested for all 2^nd^ order models by adding one of the 5 remaining variables to HD (**Fig. 3E**, middle). None of these models provided a significant increase in fit quality compared to HD alone. Therefore, the model selection procedure was terminated, and the HD model was selected as the best fitting. The full model including all 6 variables (shown for reference in **Fig. 3E**, right) did not provide a significantly better fit than the HD model alone.

To quantify tuning strength, we developed a ‘normalized tuning amplitude (NTA)’, which is equal to the curve’s trough to peak amplitude divided by the peak firing rate (i.e. the maximum value across the ‘reconstructed’ tuning curves of all variables). For this particular example cell, firing rate varied from practically zero (1 spk/s) to a peak of (104 spk/s), resulting in a NTA of 0.99.

#### Hippocampal place (AP) cell

For an example hippocampal cell that responded to AP and was modulated by ΘP, the model selection process is illustrated in **Fig. 4A**. Here, adding ΘP to the best 1^st^ order model (AP) increased the fitting quality significantly. None of the 3^rd^ order models that included AP and ΘP provided a significantly better fit than the AP+ΘP model, which was therefore selected as a best-fitting model.

The cell’s AP and ΘP response properties are summarized in **Fig. 4B,C**. When the animal explored its environment randomly, neuronal firing occurred preferentially in the lower right portion of the arena (**Fig. 4B**, left). An AP tuning curve, computed by following the usual approach of binning neuronal spiking, exhibited a place field at the corresponding position (**Fig. 4B**, middle). The fitted tuning curve constructed by the LN model resembled the experimental curve (**Fig. 4B**, left; note that is it smoother due to the LN model’s smoothness constraint; see Methods). The cell’s firing decreased from a peak 10 spk/s to a minimum of 1 spk/s, and accordingly the curve’s NTA was 0.9. The cell was phase-locked with the LFP, as shown by the fitted ΘP tuning curve (**Fig. 4C**) with a NTA of 0.57. The simulated firing rate (**Fig. 4D**, red), computed based on the AP and ΘP tuning curve, followed the cell’s measured firing well (**Fig. 4D**, black).

#### Egocentric boundary cell in RSC

The LN model incorporated a variable defined as the egocentric position of the arena’s boundary (EB). Because the arena used in the present experiments is circular, knowing the egocentric location of the entire boundary is equivalent to knowing the egocentric location of its closest point (**Fig. 5A,B**), which may be positioned anywhere within 25 cm of the head (**Fig. 5B**; see **Suppl. Fig. 1** for details). How the LN model fits the responses of an example RSC cell with EB tuning is shown in **Fig. 5C,D** (see also **Suppl. Movie 1**). This cell was largely multimodal (**Fig. 5C**): its firing rate (**Fig. 5D**) was affected significantly by head direction and linear as well as angular speed (HD, EB, LS, **Fig. 5E**), in addition to EB (**Fig. 5F**). Nevertheless, the EB response was by far the most predominant (NTA=0.96 versus NTA<=0.17 for all other significant variables).

The preferred EB response occurred in close proximity to the arena’s boundary *while* facing directly towards it, i.e. when the boundary was directly in front of the head (**Fig. 5F**, left; **Suppl. Movie** 1). In contrast, the cell was virtually silent when the animal faced away from the boundary, even when it was in close proximity (**Fig. 5F**, left; **Suppl. Movie 1**). Furthermore, the firing was independent of the animal’s allocentric location, i.e. the cell could fire anywhere along the boundary, as long as the animal was facing it. Both the experimental and fitted tuning curves exhibited a sharp peak at the corresponding location (**Fig. 5F**, middle and right panels). Note that, due to its EB tuning, the cell fired more on average when the animal’s allocentric position (AP) was close to the arena’s boundary (**Fig. 5G**). This is apparent on the *experimental* AP tuning curve, which is computed directly from experimental data (**Fig. 5G, middle**). Nevertheless, the model revealed that the cell was not sensitive to allocentric position, and the *fitted* AP tuning curve was flat.

### Overview of population responses

#### Anterior thalamic nuclei

We targeted the antero-dorsal nuclei of the thalamus in 7 animals (**Fig. 6**). Histological localization of the electrode tracks (**Suppl. Fig. 2A**) confirmed that most recordings sites were indeed located in these nuclei. Yet, we can’t exclude that some tetrodes may have contacted neighboring nuclei (e.g. antero-ventral or latero-dorsal). Therefore, we refer to these recorded regions as anterior thalamic nuclei (ATN). The proportions of cells characterized as predominantly APC, EBC, HDC, LSC and ASC are shown in **Fig. 6A**. Non-spatially modulated cells are represented by the white area of the chart, and the population of ΘP-modulated cells is outlined in black in each sector. The proportions of cells significantly modulated by any one of the 7 variables are shown as a histogram in **Fig. 6B**. Each bar is broken down into colored segments that represent the predominant variable (e.g., the variable with the highest NTA). Population responses recorded in individual animals are shown in **Suppl. Fig. 2**.

**Figure 6:**
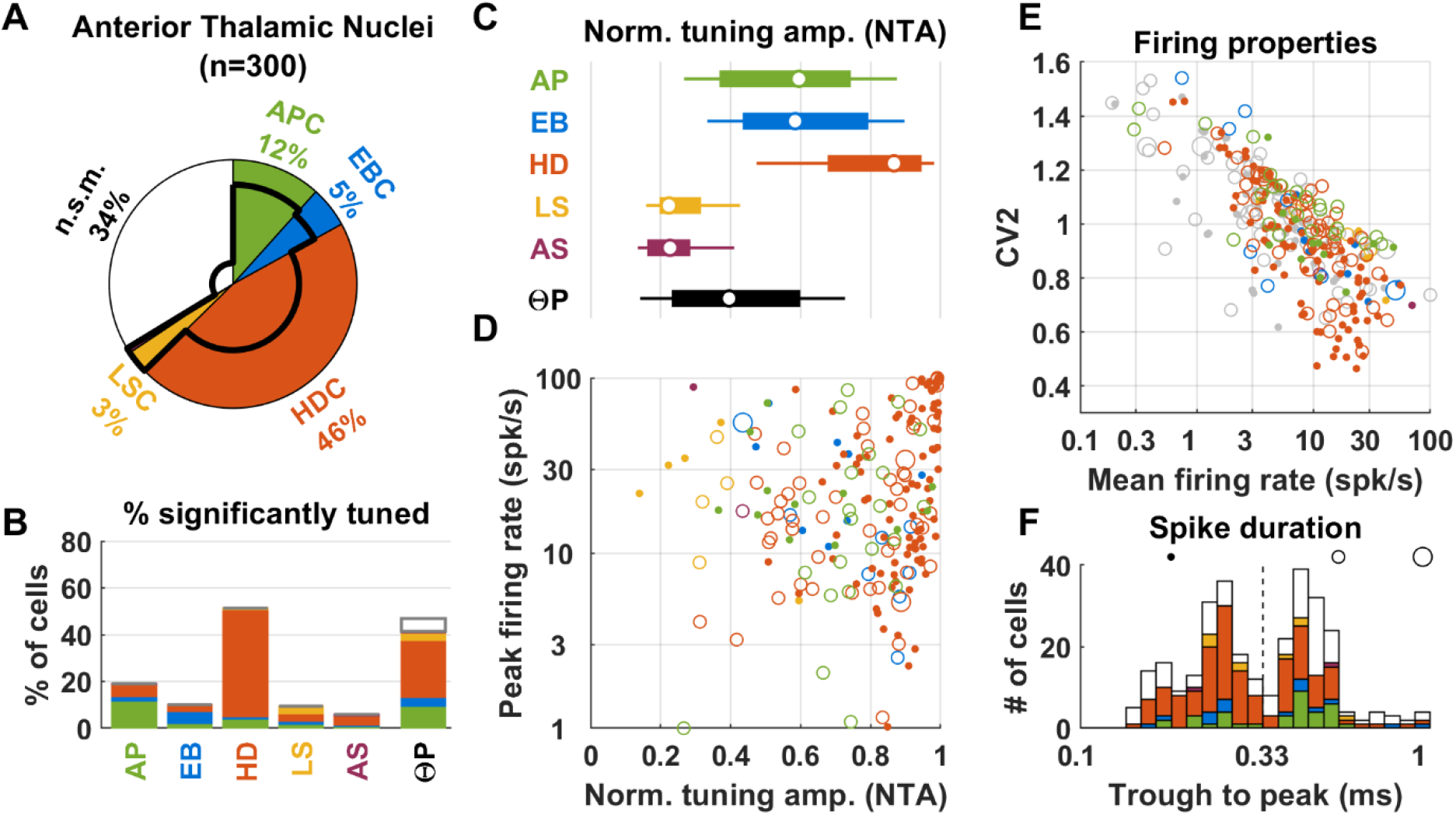
Population responses in the ATN. **A:** Diagram of cell classification. The proportions of APC, ABC, HDC, LSC and ASC are shown by colored sectors. Non-spatially modulated (n.s.m.) cells are represented by the white sector. The proportion of ΘP-modulated cells in each sector is represented by a black line. **B:** Proportion significant tuning to each variable, color-coded by cell classification based on its preferred stimulus. The white portion of the last bar represents the proportion of ΘP-modulated cells non-responsive to any other variable. **C:** Normalized response amplitude (NRA) of AP, EB, HD, LS, AS and ΘP modulation. Circles/bars/whiskers represent the median, upper/lower quartiles and upper/lower deciles. **D:** Peak firing rate versus NRA of all spatially-tuned cells (see panels A for color code). The size of the symbols encodes the trough to peak spike duration (small symbols: less than 0.33ms; medium symbols: between 0.33 and 0.8ms; large symbols: more than 0.8ms). **E:** Firing properties (CV2 versus mean firing rate), color-coded as in panel D. Not spatially modulated cells are shown in grey. **F:** Distribution of trough to peak spike duration, color-coded by cell classification. The black circle/disks illustrate the symbol size code used in panels D,E.

As expected, we found that approximately half (46%) - of ATN cells are HDC, that is, HD is the variable that modulates the cell’s firing rate the strongest (**Fig. 6A**). A small fraction of cells exhibited stronger selectivity to another variable, typically AP (12%), EB (5%) or LS (3%) (**Fig. 6A,B**, green, blue and yellow bars). Across the population, the total fraction of HD-tuned ATN cells was 51% (**Fig. 6B**). HD responses typically had very high NTA (**Fig. 6C**, orange; median across all cells with significant HD tuning = 0.87, 1^st^-9^th^ deciles: 0.47-0.98).

The peak firing rate of HDC varied widely (1^st^-9^th^ decile: 6-85 spk/s; median: 20 spk/s; **Fig. 6D**). In particular, a large cluster of HDC located in the upper right corner of **Fig. 6D** exhibited vigorous and specific firing (peak >30 spikes/s; NTA≈1), which is generally associated with “typical” HDC. Nevertheless, we also encountered a number of ATN HDC with lower peak responses.

The second spatial variable represented in the ATN was allocentric position; 19% of ATN cells had significant AP responses; **Fig. 6B**), and NTA could reach high values (median = 0.6, 1^st^-9^th^ deciles: 0.27-0.88, **Fig. 6C**, see example cells in **Suppl. Fig. 3**). A smaller proportion (10%) of ATN cells encoded egocentric boundary (**Fig. 6B**), with substantial NTA (median = 0.58; **Fig. 6C**), and 5% were classified as EBC (**Fig. 6A**). A small proportion (9%) of ATN cells were tuned to linear speed, and 3% were identified as LSC (**Fig. 6A,B**, orange). Nevertheless, linear speed responsiveness was modest (median NTA=0.22; **Fig. 6C**). Angular speed responsiveness was rare (6%) in ATN, and only 2 cells (<1%) were predominantly tuned to AS (**Fig. 6A-C**, violet).

About half (47%) of ATN cells were modulated by ΘP. Theta phase modulation was most prominent among APC, EBC and LSC (78%), compared to HDC (53%) (**Fig. 6A**). Despite the presence of a theta rhythm in all animals (**Suppl. Fig. 2**), there was a marked inter-animal variability in the fraction of ΘP-modulated HDC: almost all spatially-modulated cells in animals H51M, H54M and I29M, but almost none in animals H71M, H72M and I10M3.

APC and EBC exhibited comparable range of firing as HDC: APC: median = 18 spk/s, 1^st^-9^th^ deciles: 6-68; EBC: median = 15 spk/s, 1^st^-9^th^ deciles: 6-54. Thus, in summary, the 3 main classes of spatially tuned ATN neurons (HDC, APC and EBC) could exhibit specific responses (e.g. NTA>0.6; **Fig. 6D**) with occasionally vigorous peak responses (e.g. close to 100 spk/s, **Fig. 6D**), although lower peak responses could also be encountered (e.g. <10spk/s, **Fig. 6D**).

We also quantified additional response parameters. The neuronal firing properties (mean firing rate, CV2 and spike duration) of ATN neurons were broadly distributed (**Fig. 6E,F**): average firing rate ranged from 1.5 to 27 spk/s (1^st^-9^th^ decile; median=7 spk/s) and CV2 from 0.7 to 1.27 (1^st^-9^th^ decile; median 0.98). Firing rate and CV2 were inversely correlated (**Fig. 6E**; Spearman rank correlation=-0.75, p<10^−10^). The trough to peak duration of action potentials followed a bimodal distribution (**Fig. 6F**), thus neurons could be separated in clearly distinct groups of short-duration (trough to peak <=0.33ms, 49%) and long-duration (trough to peak >0.33ms, 51%) spikes. Neurons with short and long spike duration had similar mean firing rate (7 vs 7 spk/s, p=0.3, Wilcoxon rank test) and slightly different CV2 (0.94 vs 1.02, p=10^−3^). A further examination (**Suppl. Fig. 2**) revealed inter-subject differences in firing properties: cells in animals H51M, H54M and H59M were generally low-firing and more irregular and were distributed between short and long-duration spikes, whereas cells in animals H71M, H72M and I10M3 had higher firing rates, were more regular and had predominantly low spike duration.

Note that we could not identify any differences in variable coding among narrow and broad spiking neurons. HD cells could exhibit short or long-duration spikes (short-vs long-duration: 86 vs 51 cells, i.e. 63% vs 37%). The two groups exhibited similar mean firing rate (median: 7.3 vs 9.3 spk/s, p=0.6, Wilcoxon rank test; **Fig. 6E**) and CV2 (median: 0.92 vs 1, p=0.07). However, HDC with short-duration spikes had larger NTA (median = 0.92 vs 0.81, p=3.10^−5^) although similar peak firing (23 vs 17 spk/s, p=0.24). We also tested whether narrow and broad spiking neurons were most likely to be ΘP-modulated. To avoid a confounding factor due to inter-animal variability (where neurons H71M, H72M and I10M3 are rarely ΘP-modulated and generally narrow-spiking), we limited this analysis to H51M, H54M, H59M and I29M, and excluded non-spatially modulated neurons. We found that narrow and broad spiking neurons were equally likely to be ΘP-modulated (Chi-Square test, χ^2^=0.36, n= 1 d.o.f, p=0.54).

#### Additional ATN recordings (Peyrache et al. 2015)

In order to corroborate our finding that populations of neurons in the ATN encode AP, we analyzed previous recordings published in Peyrache et al. 2015 (**Suppl. Fig. 4,5**). Population responses (**Suppl. Fig. 4**) were similar as in our recordings, and in particular we found substantial fractions of APC in 2 out of 6 animals (**Suppl. Fig. 5**). Peyrache et al. (2015) used probe arrays (8 shanks, 200 μm spacing), which allowed us to investigate the spatial distribution of APC and HDC (**Suppl. Fig. 5**). We found that APC tend to be distributed more laterally than HDC, although the two populations overlap (**Suppl. Fig. 5C**). Furthermore, HDC and APC cells recorded in Peyrache et al. 2015 generally cover three probe shanks (**Suppl. Fig. 5A,B**), which span 400 μm laterally. Since the extent of the antero-dorsal nuclei is at most 400 μm, it is unlikely that all 3 shanks were in this nucleus; and the most lateral of the 3 shanks may have been in an adjacent nucleus (antero-ventral or latero-dorsal). In our study, recordings that identified APC in the ATN were restricted to a single track (since we used one tetrode bundle) and a narrow range of depth (**Suppl. Fig. 5I-K**), and histology indicated that tetrode tracks were located in the antero-dorsal nuclei (**Suppl. Fig. 3A**). Collectively these results suggest that APC exist in at least some portions of the antero-dorsal nuclei, although they may be more numerous in the adjacent antero-ventral or latero-dorsal nuclei.

#### Retrosplenial cortex

We implanted four animals for recording in the RSC (**Fig. 7**). One implantation (animal AA2) reached the granular cortex, where 119/180 neurons were recorded (**Fig. 7**). The other implantations (animal AA1, AA18, AA20) reached the dysgranular cortex. Responses from these regions were similar (**Suppl. Fig. 6**), thus pooled in **Fig. 7**.

**Figure 7:**
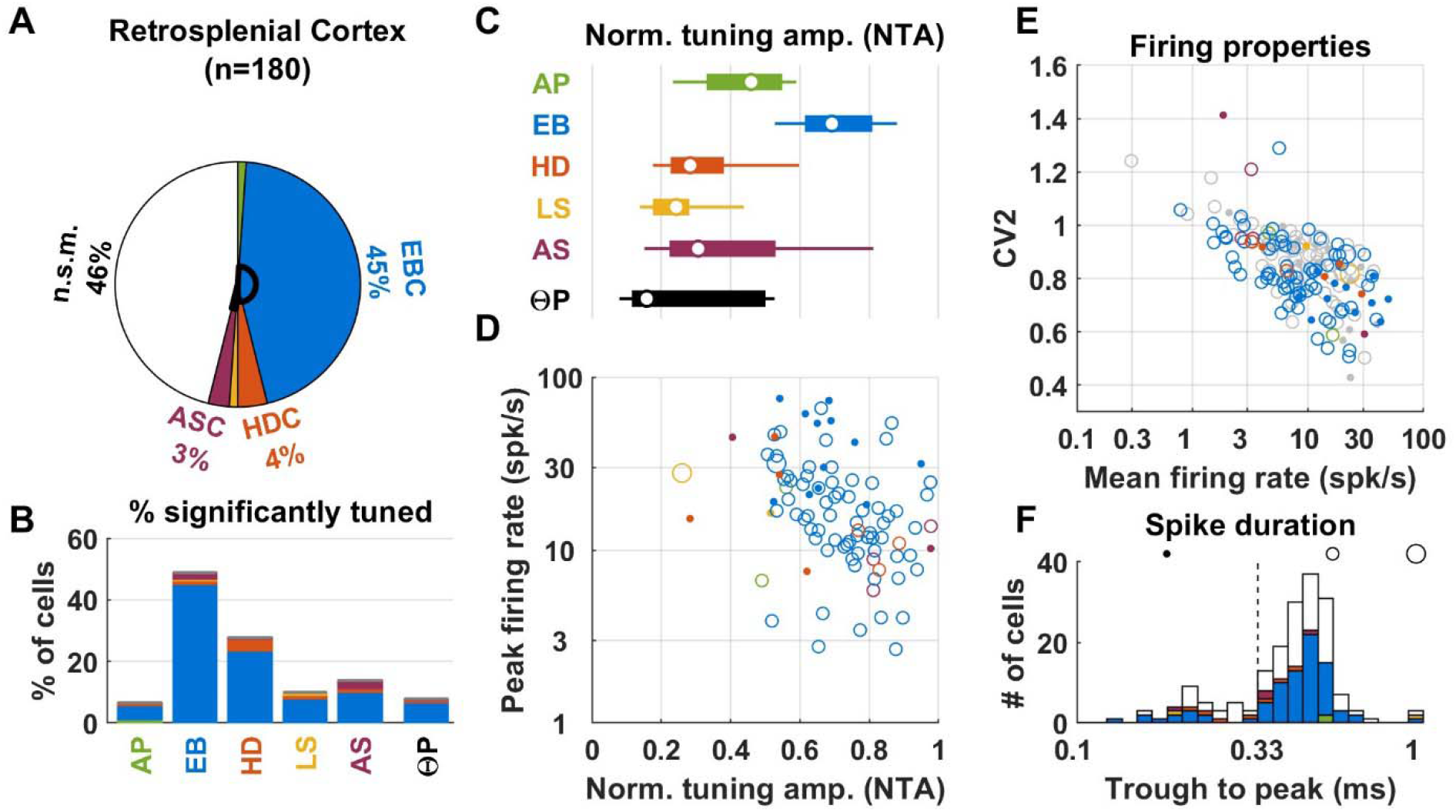
Population response in the RSC. Same legend as in **Fig. 6**.

By far, the variable represented in RSC the most was egocentric boundary (see Alexander et al. 2019). About half (45%) of RSC cells were classified as EBC. EB responses exhibited large NTA (median = 0.7; 1^st^-9^th^ decile: 0.53-0.88; **Fig. 7A-C**; an example EBC is shown in **Fig. 5** and **Suppl. Movie** 1). Peak firing rates (**Fig. 7D**) were clustered around a median value of 7 spk/s (1^st^-9^th^ decile: 17-48). We found that the ‘preferred boundary position’ of EBC (i.e. the egocentric boundary position at which their firing was maximal **Suppl. Fig. 7A**) is generally located close to the head (median = 4.95 cm, CI = [4.03-5.8]; **Suppl. Fig. 7B**). In contrast, the egocentric bearing was distributed uniformly (**Suppl. Fig. 7C**), indicating that EBC could respond when the boundary was in front of the head (**Suppl. Fig. 7A,C**: Front; F), to the right, left (R/L in **Suppl. Fig. 7C**), or behind the head (**Suppl. Fig. 7A,C**: Behind; B).

A sizeable fraction of EBC (52%) were significantly tuned to HD (**Fig. 7B**; blue portion of the bar ‘HD’), but with modest NTA (median = 0.27, 1^st^-9^th^ decile: 0.13-0.41; **Fig. 7B**). Likewise, 22% of EBC were tuned to AS (median NTA = 0.27; 1^st^-9^th^ decile: 0.14-0.48) and 15% were tuned to LS (median NTA = 0.25; 1^st^-9^th^ decile: 0.08-0.5). This indicates that EBC were often multimodal. It is striking that the majority of cells that exhibited significant HD tuning were in fact EBC.

Beyond EBC, only small fractions of responsive cells were encountered. Only 4% of the population was classified as HDC (**Fig. 7A**), although, as mentioned above, many EBC exhibited significant HD responses. Interestingly, a few (n=5, 3%) ASC were identified, and these cells had large NTA (higher than 0.8 in 4 cells; **Fig. 7D**, magenta). LFP recorded in the RSC exhibited a clear theta rhythm (**Suppl. Fig. 6**). Yet, ΘP responses were practically non-existent.

In general, RSC cells fired in a homogenous and tightly clustered fashion (**Fig. 7E**): the average firing rate was distributed around a median of 8.3 spk/s and ranged from 2.7 to 26 spk/s (1^st^-9^th^ decile). The CV2 was distributed around a median of 0.84 and ranged from 0.65 to 0.98. Most neurons (145/180, 81%) had long-duration spikes (**Fig. 7F**).

#### Cingulum fiber bundle

We recorded neuronal activity from the cingulum bundle (**Suppl. Fig. 8,9**). Most recordings (all animals except AA0; **Suppl. Fig. 9**) were located near Bregma, i.e. at the level of the transition between cingular and retrosplenial cortex along the antero-posterior axis. At this level, the cingulum conveys projections from the anterior thalamus and RSC. Therefore, we hypothesized that we would record a majority of units with short-duration spikes, consistent with axonal spikes (Barry 2015), whose firing and response properties resembled a mixture of ATN and RSC neurons. In agreement with this hypothesis, 23% of cells were classified as HDC (**Suppl. Fig. 8A,B**) and 18% as EBC (**Suppl. Fig. 8A,B**). As expected, the majority of neurons (300/380, 79%) had short-duration spikes (**Suppl. Fig. 8F**). These results show that tetrode recording from the cingulum fibers bundles are possible (and, in fact, remarkably easy); and that the section of cingulum we recorded likely conveys fibers projecting from the ATN and RSC.

#### Postsubiculum

We also analyzed previously published data recorded by Peyrache et al. (2015) in the postsubiculum of three mice (**Suppl. Fig. 10**). We found a large fraction (31%) of HDC (**Suppl. Fig. 10A**), with low or moderate firing rates (**Suppl. Fig. 10D,E**) and long spike duration (**Suppl. Fig. 10F**), consistent with layer 3 pyramidal neurons (Tukker et al. 2015, Preston-Ferrer et al. 2016, Simonnet et al, 2017; Simonnet et Fricker, 2017). The second most prominent population were APC (14%).

#### Classification of HDC: comparison with other techniques

To better appreciate how the LN model compares to classification methods used in previous studies, we evaluated HD tuning in ATN, RSC and cingulum using conventional approaches (**Suppl. Fig. 11**), which classify cells as HD-tuned if the mean vector length |R| of their experimental HD tuning curve pass a shuffling test or an arbitrary threshold (e.g. 0.26, Jacob et al. 2017, or 0.4, Yoder et al. 2009; Kornienko et al. 2018). We found that strongly tuned HD cells that pass a threshold of |R| >=0.26 are equally well detected by the LN test and other methods. We also found that a shuffling test is more sensitive than the LN model for detecting cells with moderate HD tuning (**Suppl. Fig. 11B**). However, AP or EB tuning can be erroneously interpreted as HD tuning due to sampling non-uniformity (Muller et al. 1994; Cacucci et al. 2004; Rubin et al. 2014; **Fig 7S6C**). The LN model is robust to this issue by construction, and a threshold of |R|>0.26 is high enough to rule out such cells. In contrast, using a shuffling test without accounting for AP or EB tuning may produce a moderate number of false positives, which we estimated to be 5-10% of the cells in the regions considered (**Suppl. Fig. 11G-I**). As a conclusion, the LN model is a robust technique for classifying HD cells (as well as other cell types) that agrees well with |R| tests, although it is less sensitive than shuffling tests for cells with low tuning strength.

#### Hippocampus

Many studies have focused on pyramidal place cells that are identified based on their long spike duration and typically exhibit AP responses. Here we recorded and classified the responses of a variety of cell types, including cells with short- and long-duration spikes, located mainly in the anterior part of the hippocampus.

##### AP (place) cells

A fifth (19%) of hippocampal cells were classified as APC (**Fig. 8A,B**). A significantly large fraction of APC (Chi square test, χ^2^=11, 1dof, p<10^−3^) had very long spike duration (large symbols in **Fig. 8D-F**; 10/21 APC, and 10/23 cells with very long spike duration are APC). APC with very long spike duration had strong NTA (median: 0.87, 1^st^-9^th^ decile: 0.81-0.93, **Fig. 8D**) and low peak firing (median: 5 spk/s, 1^st^-9^th^ decile: 2.7-11.4, **Fig. 8D**); while other APC had lower NTA (median 0.77, 1^st^-9^th^ decile: 0.55-0.92, p=0.012, **Fig. 8D**) and widely distributed peak firing (median 21 spk/s, 1^st^-9^th^ decile 2-56 spk/s, **Fig. 8D**).

**Figure 8:**
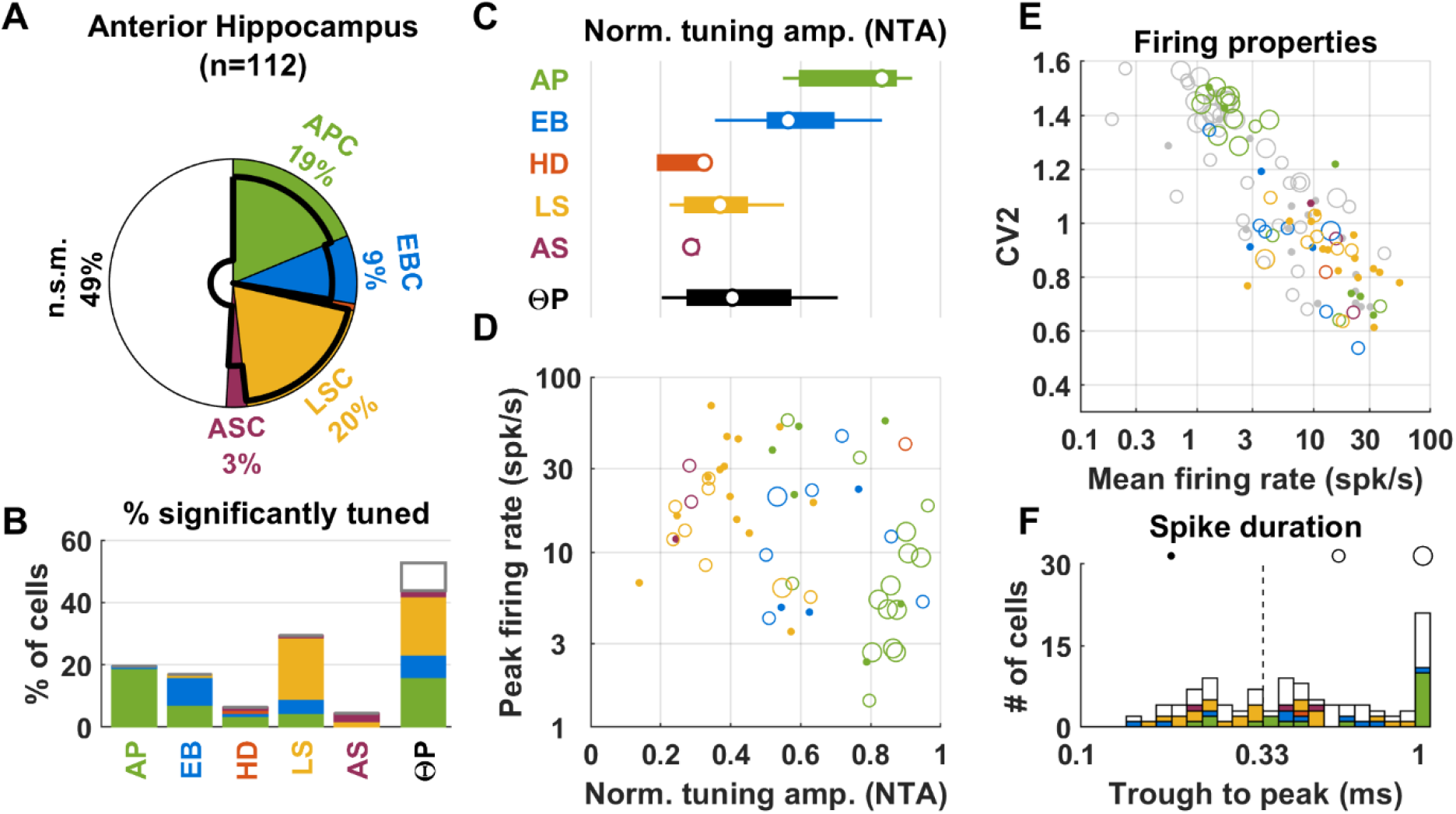
Population responses in the hippocampus. Same legend as in **Fig. 6**.

##### LS cells

One fifth (20%) of cells encoded the animal’s linear speed (**Fig. 8A**). LSC could fire short-duration (14/22, 64%) or long-duration (8/22, 36%) spikes (**Fig. 8F**). LS modulation amplitudes were modest (median: 0.37, 1^st^-9^th^ decile: 0.24-0.59, **Fig. 8C,D**).

##### Other responses

We also encountered a substantial (9%) fraction of EBC in the hippocampus **Fig. 8A,B**, with similar modulation amplitudes as in other areas (median: 0.63, 1^st^-9^th^ decile: 0.51-0.9, **Fig. 8C**).

##### Theta rhythm

Theta rhythm was recorded in all animals (**Suppl. Fig. 12**). A large fraction (53%) of all neurons was ΘP modulated, including the majority of APC (86%) and of LSC (95%) (**Fig. 8A**). Modulation amplitude was moderate (media: 0.4, 1^st^-9^th^ decile: 0.2-0.71) (**Fig. 8C**).

##### Spiking properties

As in other regions, there was a negative correlation between mean firing rate and CV2 (**Fig. 8E**, Spearman rank correlation=-0.85, p<10^−10^). Spiking duration varied widely (**Fig. 8F**): 38% of the neuronal population fired short-duration spikes (<=0.33ms trough to peak). Amongst neuron with longer-duration spikes, we observed that a large fraction (21% of the total population) fired very long-duration spikes (>0.9 ms trough to peak), and the remaining 41% fired spikes with intermediate duration. Furthermore, many neurons with >0.9ms trough to peak spike duration clustered in **Fig. 8E** (large symbols in upper left) in a group characterized by low firing rate (typically less than 4spk/s) and highly irregular (CV2>1.2).

#### Fimbria and fornix

We recorded neuronal responses in white matter regions located anterior and ventral to the hippocampus. These regions encompass the fimbria and fornix, and convey many fibers between regions of the navigation circuit (Adelmann et al. 1996; Bubb et al. 2017). Accordingly, we found that neurons recorded in these regions encode a variety of navigation variables (**Suppl. Fig. 13, Suppl. Fig. 14**).

#### Preferred phase of ΘP-modulated cells

We found that the preferred phase of ΘP-modulated cells were consistent across areas, with preferred firing occurring preferentially in the descending phase (i.e. between 180° and 270°, **Suppl. Fig. 15**).

#### Classification of APC: comparison with other techniques

We also compared the LN model to previous techniques (Skaggs 1993; Rubin et al. 2014) that compute the spatial information (SI) or AP tuning curve and use a shuffling test to evaluate statistical significance (**Suppl. Fig. 16**). Again we found that the LN model is less sensitive than the shuffling test (**Suppl. Fig. 16B**). Furthermore, we found that HD tuning could easily be misinterpreted as AP tuning, as pointed out by (Peyrache et al. 2017) (**Suppl. Fig. 16C**, left), an issue that could also affect EBC (**Suppl. Fig. 16C**, right). Because of this, 17 to 24% of the cell populations in ATN, RSC and cingulum were incorrectly classified as APC by a conventional shuffling test. Thus, a traditional measure of spatial information combined with a shuffling test may be enough to identify APC reliably in the hippocampus, where EBC and HDC are scarce (**Suppl. Fig. 16K**). In contrast, testing for AP responses in regions that host other cell types, such as the ATN, RSC, cingulum and parahippocampal regions requires classification methods that can control for responses to other variables, such as the LN model or techniques used in earlier studies (Muller et al. 1994; Cacucci et al. 2004; Rubin et al. 2014; Peyrache et al. 2017).

## Discussion

We used an LN model (Hardcastle and al. 2017) to analyze how neuronal activity in multiple areas of the brain’s navigation network encode combinations of navigation variables. As expected, the LN model identified prominent and well-known characteristics of these areas, such as the prevalence of HDC in ATN and postsubiculum, or place cells in the hippocampus, but also revealed several novel features. We found that egocentric information is represented extensively not only in RSC (Alexander et al. 2019), but also in ATN and hippocampus. Furthermore, ^~^12% of ATN neurons encode the animal’s allocentric position, a finding that we confirmed by re-analyzing previously published data (Peyrache et al. 2015). We recorded spiking activity from the cingulum fiber bundle and found that most units resemble ATN or RSC neurons, suggesting that spatial information encoded in these regions travels along the cingulum.

### Anterior thalamic nuclei

Our recordings in the ATN consisted predominantly of antero-dorsal nuclei neurons. The presence of a large population of HDC in the ATN of rats (Taube et al. 1995; Blair and Sharp 1996; Zugaro et al. 2001; Peyrache et al. 2015, 2017; Page et al. 2017) and mice (Yoder and Taube 2009) is well documented.

More surprising was the finding of a substantial population of APC, i.e. cells tuned to the animal’s allocentric position, in 3 of our animals (**Suppl. Fig. 2**) as well as in 2 animals in a previously published dataset (Peyrache et al. 2015; **Suppl. Fig. 5**), some of which exhibited very sharp tuning (**Suppl. Fig. 3**). Importantly, the spike waveform of most APC exhibited long trough to peak duration (>0.33 ms; **Fig. 6F** and **Suppl. Fig. 4F**, green), indicating that they were not likely fibers traversing the ATN. These APC responses could not have been recorded in the fimbria, just dorsal to the thalamus, and erroneously classified as ATN cells because, although the fimbria contains 12% of APC (**Suppl. Fig. 13A**), these cells typically exhibit short trough to peak duration (**Suppl. Fig. 13F**). Yet, we cannot resolve with complete certainty the location of these neurons. The dataset of Peyrache et al. (2015) indicates that these neurons may overlap the population of HDC in the antero-dorsal nuclei, but be more numerous lateral to the antero-dorsal nuclei, i.e. in the antero-ventral or latero-dorsal nuclei (**Suppl. Fig. 5**). Thus, more detailed studies would be required to assess the spatial distribution of APC in the anterior thalamus.

Theta rhythm is a fundamental property of the navigation circuit, thought to mediate memory and planning in the hippocampus (Buzsáki and Moser 2013) and in general inter-regions communication (Maris et al. 2016). Previous studies (Vertes et al. 2001, Albo et al. 2003) have shown that neurons in the ATN, including the antero-ventral and antero-dorsal nuclei, are modulated by the *hippocampal* theta rhythm. Furthermore, Tsanov and colleagues (2010, 2011) have shown that a portion of HDC in the antero-ventral nuclei of rats exhibit a rhythmic firing at theta frequency. In this study, we targeted the ATN of 7 animals, and histology (**Suppl. Fig. 2A**) indicates that most recording took place in the antero-dorsal nuclei. We found that a clear theta-band LFP was present in all animals (**Suppl. Fig. 2E**, peak in the 5-12Hz frequency range), and that over half of HDC and most APC and EBC were modulated by local theta rhythm. Collectively, these studies indicate that at least part of HDC as well as EBC in the ATN are modulated by a theta rhythm that may originate in the hippocampus. The hippocampal formation projects to the ATN directly, though projections of the subiculum to the antero-ventral nuclei and of the postsubiculum and parasubiculum to the antero-dorsal nuclei. It also projects indirectly via the mammillary nuclei (Kocsis 1997; Vertes et al. 2001; Vann and Aggleton 2004).

### Retrosplenial cortex and Egocentric boundary cells

We found that about half of RSC neurons encode the egocentric position of the arena’s boundary. The existence of egocentric boundary cells (EBC) was proposed by Derdikman (2009), and EBC were recently identified in the entorhinal cortex (Wang et al. 2018, Gofman et al. 2019), postrhinal cortex (Gofman et al. 2019), parasubiculum (Gofman et al. 2019), striatum (Hinman et al. 2019) and RSC (Alexander et al. 2019). Several studies have described HDC in the RSC (Chen et al. 1994a,b; Cho et al. 2001; Jacob et al. 2017; Lozano et al. 2017). Here we found that, although 28% of RSC cells were tuned to HD, the majority of these were in fact EBC with weaker but significant responses to HD, such that only 4% of RSC cells were classified as HDC. Over half of EBC in the RSC were significantly tuned to HD (similar to EBC in the parasubiculum and medial enthorinal cortex, Gofman et al. 2019), which suggests that the RSC may be involved in combing multiple reference frames (see Clark et al. 2018). Note that the population responses appear similar in the granular (animal AA2; **Suppl. Fig. 6**) and dysgranular (other animals; **Suppl. Fig. 6**) cortices.

### Cingulum fiber bundle

The cingulum connects several areas of the navigation system (Domesick 1970; van Groen and Vyss 1990, 1995, Bubb et al. 2017, 2018). It conveys anterior thalamic projections to the RSC and parahippocampal regions and RSC projections to the cingulate cortex and parahippocampal regions. The cingulum also carries projections from the subiculum to the RSC and parahippocampal regions, however it is unlikely that these fibers were recorded since most of our cingulum recordings were performed at the level of the anterior part of the RSC, i.e. ^~^0.2mm posterior to Bregma. Lesion studies have confirmed that the cingulum is involved in using allocentric cues for spatial navigation (see Bubb et al. 2018 for review), suggesting that it conveys spatial information.

The present recordings from the cingulum identified large fractions of HDC and EBC that are strikingly similar to their ATN and RSC counterparts. In particular, many HDC in the cingulum exhibited the high firing rate and NTA (**Suppl. Fig. 9D**) that are typical of ATN HDC (**Fig. 6D**) and would scantly be distinguishable from ATN units when examined online. Likewise, EBC recorded in the cingulum were similar to those encountered in RSC (**Fig. 7D, Suppl. Fig. 9D**), but as expected exhibited shorter waveforms (Robbins, 2013).

### Hippocampus

As expected, we found a population of APC in the hippocampus that matched the typical profile of hippocampal place cells in rats (O’Keefe 1971, 1796) and mice (Muzzio, 2009; Jeantet, 2012; Kinsky, 2018), i.e. high spatial selectivity, low average firing, and long spiking duration (**Fig. 8D-F**). We also recorded a population of cells that were tuned to linear speed. In agreement with a recent study (Góis and Tort 2018), these cells often exhibited fast spiking and short spike durations.

### Fimbria and fornix

Neural activity in the white matter regions located anterior and ventral to the hippocampus, regions that contain a variety of fiber tracks, including projections from the septal nuclei and entorhinal cortex as well as commissural projections (Adelmann et al. 1996; Bubb et al. 2017), showed a variety of spatially modulated signals. In particular, a sizable population (12%) of APC was identified (**Suppl. Fig. 13A**), although its properties were clearly distinct from those of typical hippocampal place cells (**Fig 8A**). Most notable is the fact that 16% of recorded cells were HDC (**Suppl. Fig. 13A**) with large firing rate and NTA (**Suppl. Fig. 13D**). In many animals (e.g. H71M, I29M, VR8, **Suppl. Fig. 14**), these HDC were recorded directly above the thalamus. This reveals a methodological challenge for targeting ATN, since it shows that lowering electrodes until characteristic HDC are observed is not sufficient because similar response properties are common above the thalamus. Although tetrodes are rarely used to target fibers, previous studies (Robbins et al. 2013) have shown that they can be used to record axonal spikes. We found that spiking could readily (and, in fact, remarkably easily) be recorded in the fiber tracks. Although fiber tracks traveling in the fimbria are difficult to identify, the cingulum is anatomically well delimited, and our study demonstrates that systematic investigations of information transmitted along this bundle is feasible.

In summary, the LN model (Harcastle et al. 2018) offers a more robust way to investigate neurons responses recorded in areas of the brain’s navigation system by allowing to test for multiple variables at once while proving remarkably immune to pitfalls such as overfitting. In contrast, traditional methods of computing HD tuning curves or place fields often proved impracticable when applied across all brain areas (Suppl. Fig. 8, Suppl. Fig. 11): for instance, a traditional place field analysis designed for hippocampal place cells produces aberrant results in the thalamus because it is biased by HD tuning (Suppl. Fig. 11). Our study provides a new picture of the information encoded by ATN and RSC cells as well as their output bundle, while re-visiting the responses in the anterior hippocampus and the neighboring white matter areas, and may serve as a guide for future investigations in these areas.

## Methods

### Animals

A total of 22 male adult mice (21 C57BL/6; 1 nNOS-ChR2 BAC C57BL/6 transgenic mouse: VR8), 3-6 months old, were used in this study. We implanted a head-restraining bar and a microdrive/tetrode assembly under general anesthesia (Isoflurane) and stereotaxic guidance. Animals were single-housed on a reversed [12/12] light/dark cycle. Experimental procedures were conducted in accordance with US National Institutes of Health guidelines and approved by the Animal Studies and Use Committee at Baylor College of Medicine (protocol n°AN-5995).

### Neuronal recordings

Neurons were recorded using 6 (animals AA1/AA2/AA18/AA20), 5 (animal VR8) or 4 (all other animals) tetrode bundles constructed with platinium-iridium wires (17 micrometers diameter, polyimide-insulated, California Fine Wire Co, USA) and platinum-plated for a target impedance of 200kΩ using a Nano-Z (Neuralynx, Inc) electrode plater. Tetrodes were cemented to a guide tube (26-gauge stainless steel) and connected to a linear EIB (Neuralynx EIB/36/PTB). The tetrode and guide tube were attached to the shuttle of a screw microdrive (Axona Ltd, St Albans, UK) allowing a travel length of ^~^5mm into the brain.

Tetrodes were positioned under stereotaxic guidance. We targeted the ATN by implanting 0.2mm posterior and 0.7mm lateral relative to Bregma, and placing the tetrodes at an initial depth of 1.8mm relative to the surface of the cortex. The cingulum was targeted by implanting at the same coordinates but at an initial depth of 1.2mm (except for one animal, AA0, where the cingulum was reached 2mm posterior to Bregma). The anterior hippocampus was targeted by implanting 0.6mm posterior and 0.7mm lateral relative to Bregma. Recordings in the fimbria were obtained along the track of electrodes targeting the hippocampus, ATN or cingulum. The RSC was targeted by implanting 2mm posterior and 0.07mm (AA2/AA18), 0.5mm (AA20) or 0.7mm (AA1) lateral relative to Bregma and placing the electrodes at the surface of the cortex.

LFP were recorded as a low-frequency content of the neuronal data, referenced to a ground screw implanted in the skull. Similar to Harcastle et al. 2015, we downsampled LFP data to 250Hz and band-pass filtered it in the 4Hz-12Hz range (second-order Butterworth filter).

At the end of the study, the animals underwent transcardial perfusion with 4% paraformaldehyde (PFA). The brains were postfixed in 4% PFA and then transferred to 30% sucrose overnight. Brain sections (40μm) were stained (Nissl or neutral red staining), and examined using bright-field microscopy to localize tetrode tracks.

### Recording procedure

We recorded neural activity while animals explored a circular arena (50cm diameter, 30cm height). Recordings were performed in 8-minutes sessions. The walls of the arena where white, with a black cue card covering an angle of 45°. Illumination was provided by a LED strip lining the top of the white section of the arena’s inner wall. The tetrodes were connected to a tethered head stage that included two LEDs (1 red and one infra-red, 4 cm apart) for optical tracking (Cineplex 3, Plexon Inc.). Broadband neuronal data were acquired at 22 kHz using a MAP system (Plexon Inc., Rasputin V2 software) and stored for offline analysis. Spike sorting was performed manually based on trough and peak spike amplitude and principal component analysis, using a custom Matlab script. Data was stored on a custom database programmed using Datajoint (Yatsenko et al. 2015).

### Data analysis

We used optical head tracking data to compute the following variables, which were divided into bins to fit the LN model: (1) Allocentric position of the head (AP) in 2D, which ranged from −25 to 25cm in each dimension, and was binned using a 12×12 grid. Note that AP is always included in a 25cm radius circle, since the arena was circular, and therefore the corners of the 12×12 were never covered. However, the LN model is designed in such a way that adding ‘empty bins’ won’t affect its results. (2) Egocentric position of the arena boundary (EB) was encoded in 2D, as described in **Suppl. Fig. 1**. Similar to AP, EB ranged from −25 to 25cm, is always included in a 25cm radius circle, and was binned using a grid defined in **Suppl. Fig. 1**. (3) Head direction (HD), which ranged from −180 to 180°. A HD of 0° corresponds to the direction of the center of the black cue card. HD was divided in 18 bins. (4) Linear speed (LS) which ranged from 0 to 30cm/s and was divided in 18 bins. (5) Angular speed (AS) which ranged from −150 to 150°/s and was divided in 18 bins. (6) Phase of the theta-band LFP (ΘP) was computed by applying a Hilbert transform to band-pass filtered (5-12Hz) LFP signals, as in Hardcastle et al. 2017. ΘP ranged from 0 to 360° and was divided in 18 bins.

We used the same forward search procedure as in Hardcastle et al. (2015) to chose the best fitting model.

Additional variables were tested but were excluded from the analysis after we determined that they didn’t contribute meaningfully to neuronal responses. We tested: (1) the egocentric direction of the center of the cue card; (2) the point of the arena’s boundary that the head faced, i.e. we computed the intersection of the head’s forward axis and of the arena boundary and divided the arena boundary in bins, and (3) the frequency and (4) the magnitude of the LFP.

To ensure that results were robust, we excluded all recordings where AP covered less than 2/3 of the bins included in the 25cm radius circle. We verified that this criterion was sufficient to ensure that all other variables were well sampled. Two animals (H62M, H65M) were excluded from the analysis because no session passed this criterion.

We computed the CV2 of the spike trains as the median value of (2. |ISI_i+1_-ISI_i_|/(ISI_i+1_+ISI_i_)) across all inter-spike intervals (ISI).

### LN model fitting

Model fitting was performed by using the Matlab code provided by the authors of (Hardcastle et al. 2017). We optimized the code to allow fitting a large number of models (up to 9 in preliminary testing). We programmed it to fit only the models that were necessary for the forward search procedure. If n variables are tested, this procedure will test, in a worst case, n 1^st^ order models, n-1 2^nd^ order models, n-2 3^rd^ order models, and so on until 1 n^th^ order model; and the procedure will generally terminate earlier. Therefore, fitting only these models instead of all possible models reduces the complexity from 2^n^-1 in all cases to n.(n+1)/2 in the worst case.

The LN model scores each model based on a log-likelihood measure. When a cell was recorded during multiple sessions, the models were fitted to each session separately, and the resulting log-likelihood averages across sessions.

Note that, when a model fits continuous firing rates as the LN model does (as opposed to firing rate averaged across several trials), its coefficient of correlation will be heavily affected by the neuron’s firing variability, especially in the case of sparsely firing neurons such as hippocampal place cells. Since the LN’s model statistical analyses are based on log-likelihood, and comparing coefficients of correlation across cell types and areas would easily be misleading, we opted not to report coefficients of correlation.

### Shuffling test for HD and AP tuning

We also quantified HD tuning by computing the mean vector length |R| of the experimental HD tuning curve. HD tuning curves were computed as a histogram FR(HD) with a bin width of 20° and smoothed using a Gaussian kernel (standard deviation 15°). |R| was computed as |R| = R = c.∑ FR(HD)*exp(-i*HD)/ ∑ FR(HD) with c = 3.6*π/180/2/sin(1.8) (Zar, 1998).

We quantified AP tuning of the experimental AP tuning curve by computing the spatial information SI. We divided the area in pixels (2.5cm width, Bjerknes et al. 2018) as computed SI = ∑p_i_.FR_i_/FR_avg_.log_2_(FR_i_/FR_avg_), where FR_i_ is the firing rate in the i^th^ pixel, FR_avg_ the average firing rate, and p_i_ the probability of being in the i^th^ pixel.

We implemented a shuffling test to assess the significance of |R| and SI. For each cell, we generated 1000 samples by circularly shifting the spikes train by a random value of at least ±10s (i.e. the shifted trial was wrapped to the beginning) and we recomputed |R| and SI. The un-shuffled values of |R| and SI were considered significant if the exceeded 99% of the shuffled values.

### Additional data from Peyrache et al. 2015, 2017

In order to supplement our recordings in the ATN, we analyzed published recordings (Peyrache et al. 2015, 2017) performed in the ATN of 6 mice (named AP12, AP17, AP20, AP24, AP25, AP28, AP32 here, corresponding to Mouse12, Mouse17, etc in the original dataset), as well as in the post-subiculum of 3 mice (AP24, AP25, AP28) while animals walked freely in a rectangular arena (53×46 cm). We included a total of 39 sessions where animals covered the arena uniformly. We selected the shanks in which most neurons were recorded, which likely correspond to the antero-dorsal nuclei and their immediate vicinity, for population analysis (see **Suppl. Fig. 4,5**).

### Statistics

We used a threshold value of 0.01 in all statistical tests. All statistical tests used in this study are non-parametric. All comparisons between median values are performed using double-tailed Wilcoxon rank tests.

## Supporting information

Model fit and firing properties of all neurons

## Data availability

We provide a Supplementary Data spreadsheet as supplementary information. This spreadsheet contains the response statistics of all cells shown in Figs. 6,7,8 and Suppls. Fig. 2,3,4,6,8,9,10,12,13,14.

## Code availability

Data was analyzed based on code provided by the authors of (Hardcastle et al. 2017). Code used specifically in this study will be made available upon request.

## Acknowledgements

Supported by the Simons Collaboration on the Global Brain Grant 542949, DC004260, DC014518 and DC015602.

---

**Supplemental Figure 1:**
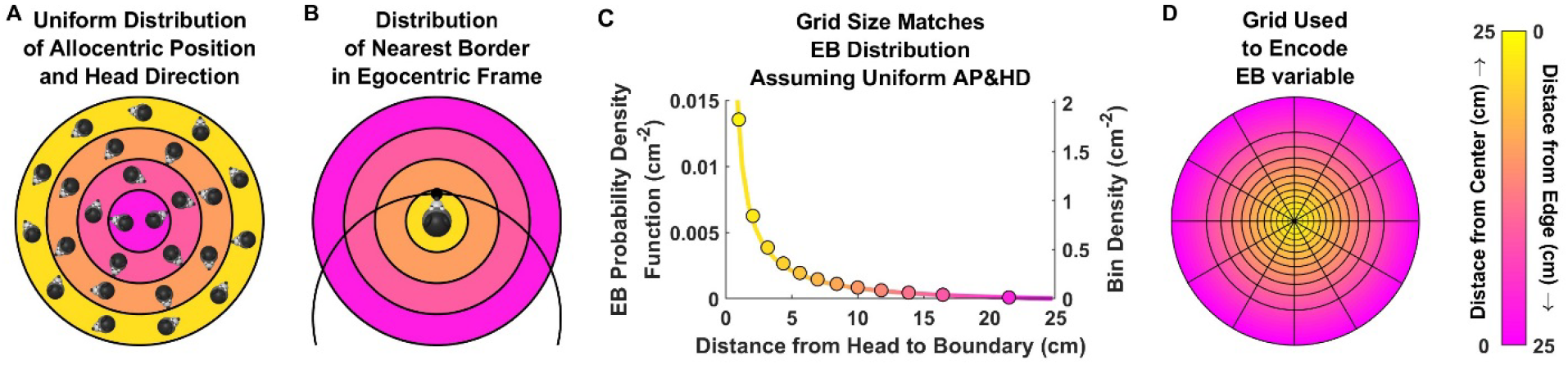
Encoding the EB variable. **A:** Arena seen from an allocentric point of view. The arena is divided in color-coded (yellow to magenta) concentric regions. All regions have the same width; but outermost regions have a larger perimeter. Therefore, the outer regions (e.g. yellow) have a larger surface area than the innermost (e.g. magenta). If the animal explores the arena uniformly (as illustrated by the mice heads, where AP and HD are uniformly distributed), the head is more likely to be located in the yellow regions. **B:** In egocentric space, the egocentric position of the nearest boundary (EB) falls inside of a circle with 25 cm radius (since the closest boundary is at most 25 cm away from the head). Points where the head is close to the boundary correspond to the (large) yellow region in A and to the (much smaller) yellow region in B. Thus, if AP is uniformly distributed in A, EB is non-uniformly distributed in B, with a higher probability of being close to the center. **C:** Probability density function of EB (line), estimated by drawing a large (10^7^) number of head positions where AP and HD are uniformly distributed (as in panel A; we assumed that AP can’t be located closer than 0.5cm to the arena’s wall to account for the animal’s head size) and computing the corresponding EB. The density function is expressed in probability per cm^2^, and plotted here as a function of the distance between the head and the boundary. The density is higher in proximal space (i.e. yellow region). Accordingly, we bin the egocentric space in B with a grid that has a higher density (i.e. small bins) in proximal space (dots, see panel **D**). **D:** To represent EB, we created a grid that has a higher resolution in proximal space so as to match the distribution in (C). First, we computed 12 concentric zones whose width was adjusted such that each contained 1/12^th^ of all samples used in C. Thus, by construction, EB is distributed uniformly across all zones. All zones were further divided into 12 angular sectors to create a grid onto which EB is uniformly distributed. The surface area of grid bins is computed and its inverse (grid density, in cm^−2^) is shown, as a function of radius, in C. As expected, the resulting points (disks) scale with the EB density function.

**Supplemental Figure 2:**
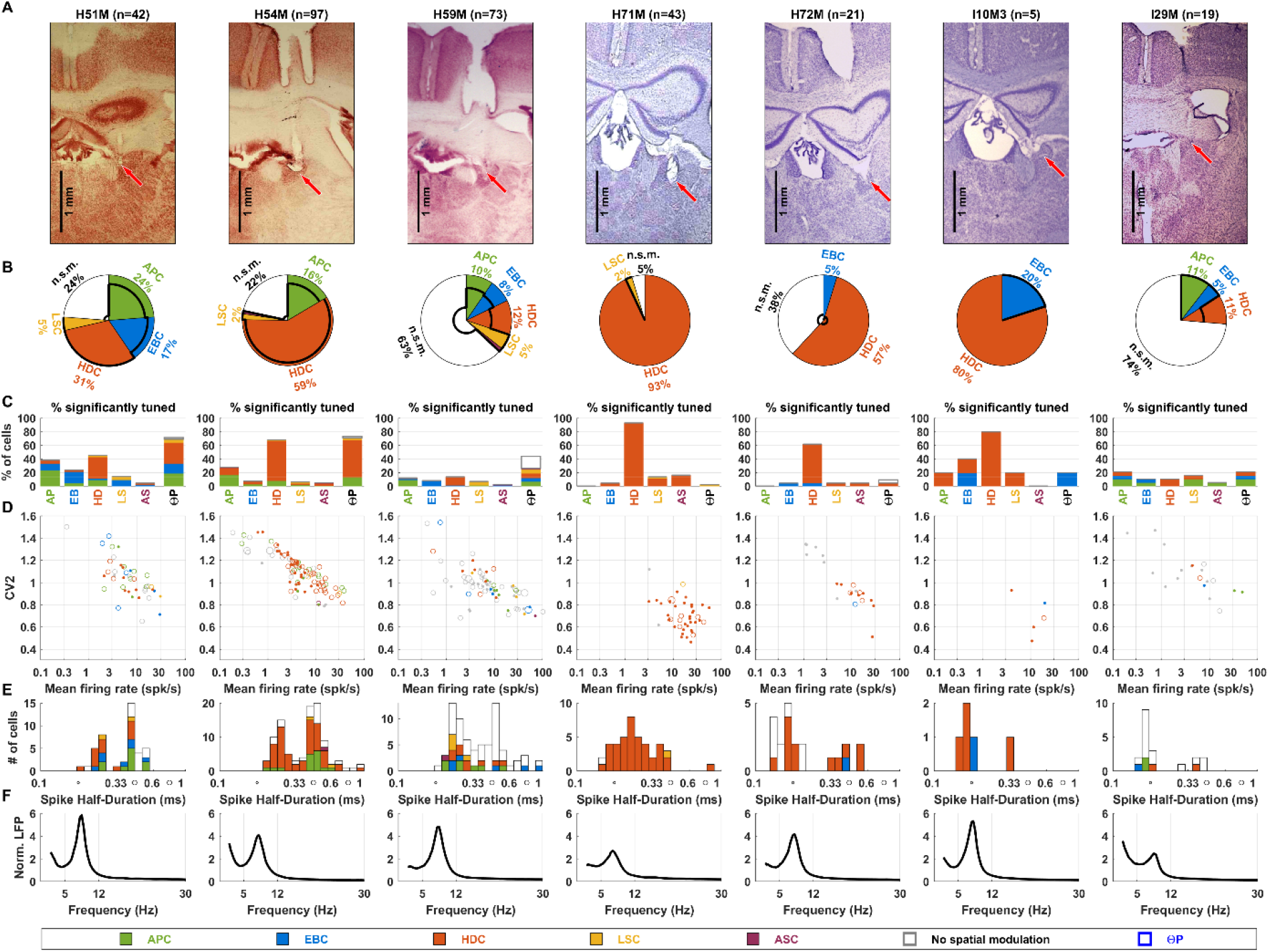
Summary of ATN properties of individual mice. **A:** Coronal histology sections in each animal, showing the location of tetrode tracks. Tracks are predominantly located in the antero-dorsal nuclei, although they may possibly have contacted neighboring regions; e.g. antero-ventral nucleus in H71M; latero-dorsal nucleus in H72M; stria medullaris in H51M. **B:** Venn diagram showing the proportions of all cell types and of ΘP-modulated cells, as in **Fig. 6A**. HDC were recorded in all animals. Clear populations (>5%) of APC and EBC were recorded in 3 and 4 animals respectively. **C:** Proportion significant tuning to all variables, as in **Fig. 6B**. **D:** Firing properties (CV2 versus mean firing rate) of recorded cells, as in **Fig. 6E**. **E:** Distribution of trough to peak spike duration, as in **Fig. 6F. F:** Average (over all electrodes and recording sessions) LFP power spectrum. A clear theta-band LFP, peaking between 5 and 12Hz, was observed in all animals.

**Supplemental Figure 3:**
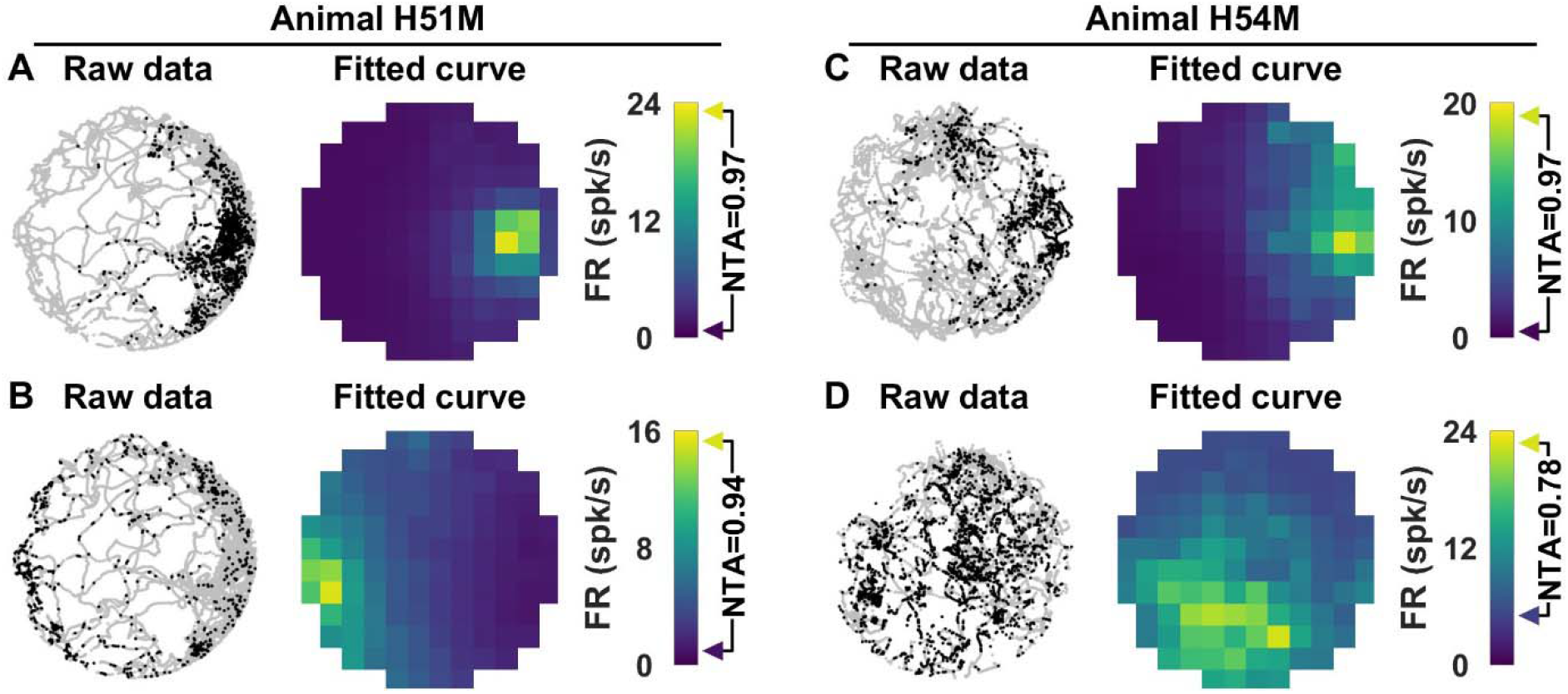
Example APC cells in the ATN. Left panels: raw data, showing the recorded head position (grey) and spikes overlaid as black dots. Right panels: fitted tuning curve represented as color maps. The NTA is indicated on the color scale.

**Supplemental Figure 4:**
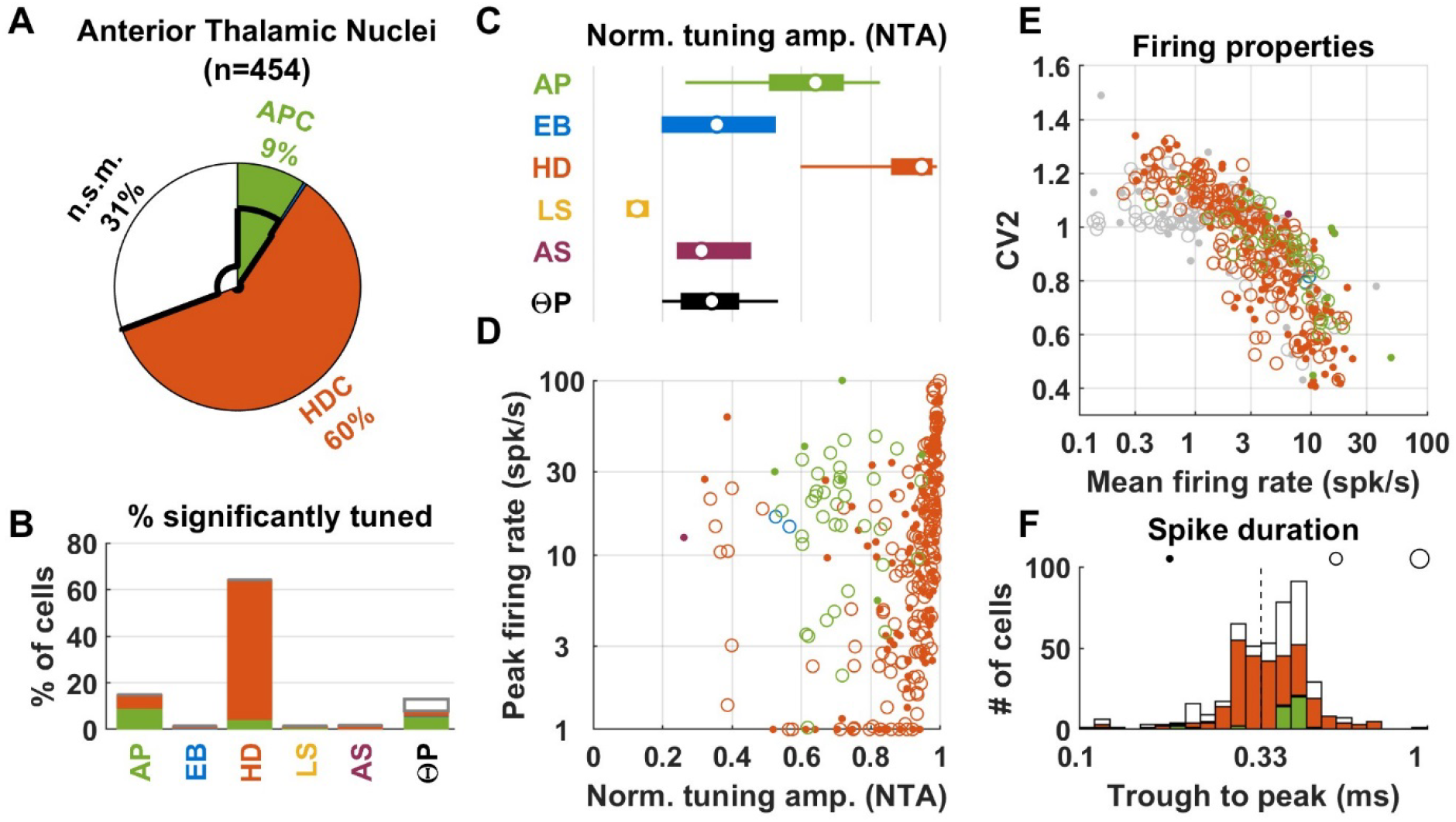
Population responses in the ATN from previously published data (Peyrache et al. 2015). Same legend as in **Fig. 6**. In agreement with our recordings, a large fraction (60%, panel A) of ATN cells are classified as HDC, with high NTA and peak firing (e.g. NTA>0.9, peak firing > 30 spk/s), similar to our recordings. Across all significantly-tuned cells, HD responses have high NTA (panel C; median = 0.95; 1^st^ – 9^th^ decile: 0.6-0.99). Note that HDC with very low peak firing (e.g. <3 spk/s) amounted to a larger fraction of the population than in our recordings (23% vs. 3%; panel D). It is possible that longer recording durations used in Peyrache et al. 2015 (30 min foraging, total recordings amounting to several hours) made it easier to identify clusters of spikes occurring at low frequencies. In agreement with our recordings, 9% of cells were classified as APC (panel A; see also **Suppl. Fig. 5**); median NTA = 0.64; 1^st^ – 9^th^ decile: 0.27-0.83; current study: = 0.63; 1^st^ – 9^th^ decile: 0.26-0.88; Wilcoxon rank sum test: p=0.99). Likewise, the distribution of APC’s peak firing rates (median = 20 spk/s; 1^st^ – 9^th^ decile: 3.6-41) matched our recordings (median = 17 spk/s; 1^st^ – 9^th^ decile: 5-54; Wilcoxon rank sum test: p=0.99). Similar to our recordings, we found a mixture of short-duration (39%) and long-duration (61%) spikes (panel F), although the bimodality was not as pronounced as in our data (compare with **Fig. 6F**) possibly because of differences in electrode type and filtering.

**Supplemental Figure 5:**
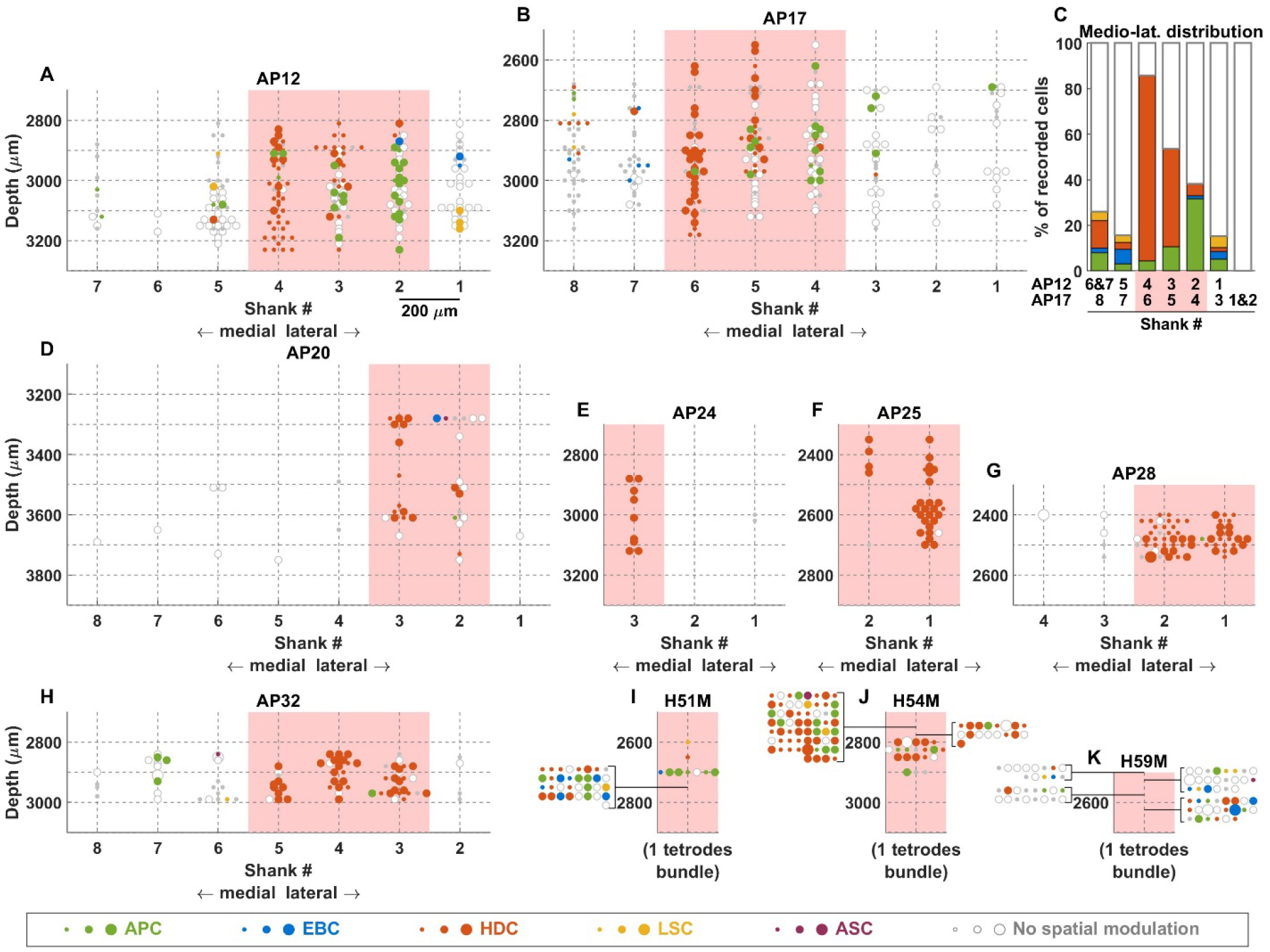
Spatial distribution of neuronal response types in ATN from previously published data (Peyrache et al. 2015) using multiple-shank probes (200 μm spacing between shanks). **A,B:** Cell location in animals AP12 and AP17, where most APC were found (abscissae: shank number, with shanks distributed along the medio-lateral axis; ordinate: depth relative to Bregma). Neurons recorded on a single shaft are staggered laterally for better visualization. Pink zone: region with most responsive neurons, likely the antero-dorsal nucleus and its immediate vicinity. Only neurons recorded in this region were included in the population analysis in **Suppl. Fig. 4**. HDC tend to cluster medially, APC laterally. **C:** Percentage of cells of each classification as a function of medio-lateral position in animals AP12 and AP17, shifted to align the presumed location of ATN (pink). HDC were predominant (82% of recorded cells) in shanks #4 (AP12) and #6 (AP17) (leftmost bar), whereas APC were found more often in shanks #2 (AP12) and #4 (AP17) (32% of recorded cells). Thus, HDC and APC are spatially segregated in ATN, although with some overlap. **D,H:** Location of neurons recorded in other animals by Peyrache et al. 2015. **I,K:** Location of neurons in the present study (one tetrode bundle), for animals where APC were present in the ATN intermingled with HDC. Large groups of cells recorded at a single location are represented at the side of the graph for readability. We observe that HDC and APC are commonly recorded at a single location. This indicates that these recordings were performed in a region where APC and HDC overlap.

**Supplemental Figure 6:**
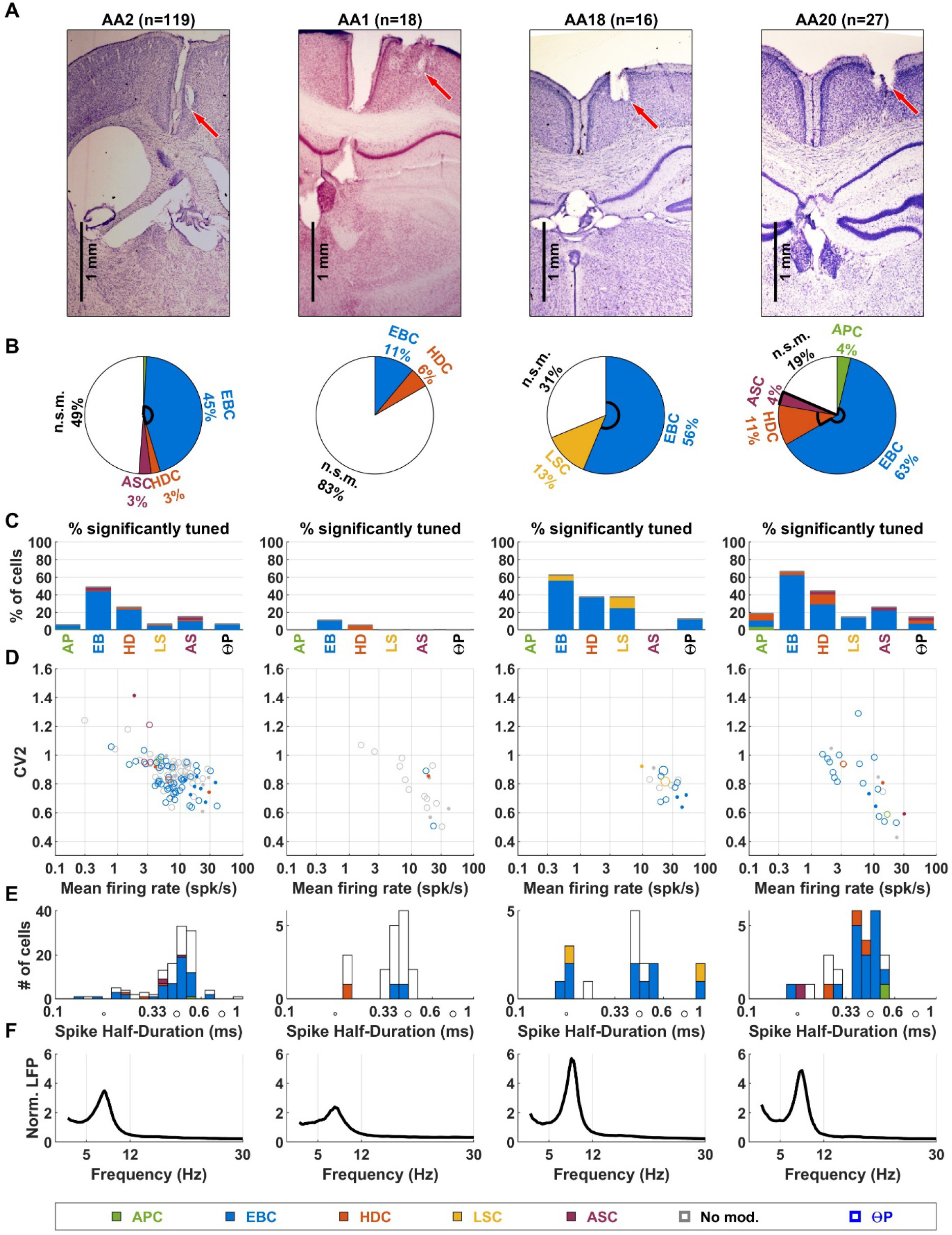
Recordings in the RSC of individual mice. Same legend as in **Suppl. Fig. 2**. Animal AA2 was implanted in the granular cortex, and AA1, AA18 and AA20 in the dysgranular cortex.

**Supplemental Figure 7:**
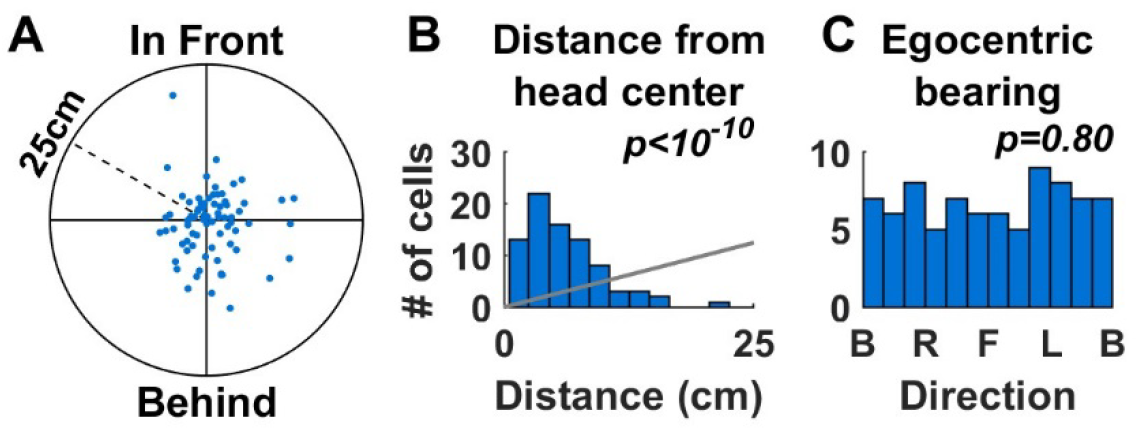
Spatial properties of EBC in RSC. **A:** Distribution of the preferred position of EBC. The preferred position refers to the egocentric location of the nearest boundary (as in **Fig. 2,5**) at which the cell fires most, and is plotted in polar coordinates, with the distance from the origin representing the distance from the head, and the direction representing the egocentric bearing to the nearest point (i.e. in front of the head, to the right, behind the head, or to the left). **B:** Distribution of the distance from the head to the preferred position (histograms). Distribution expected if points were distributed uniformly in panel A are shown in gray (H_0_). P-values are computed based on Kolmogorov-Smirnov test. **C:** Distribution of the egocentric bearing to the preferred position. B, R, F, L refer to preferred position occurring behind, right, in front or left of the head. P-values are computed based on circular Rayleigh test for uniformity.

**Supplemental Figure 8:**
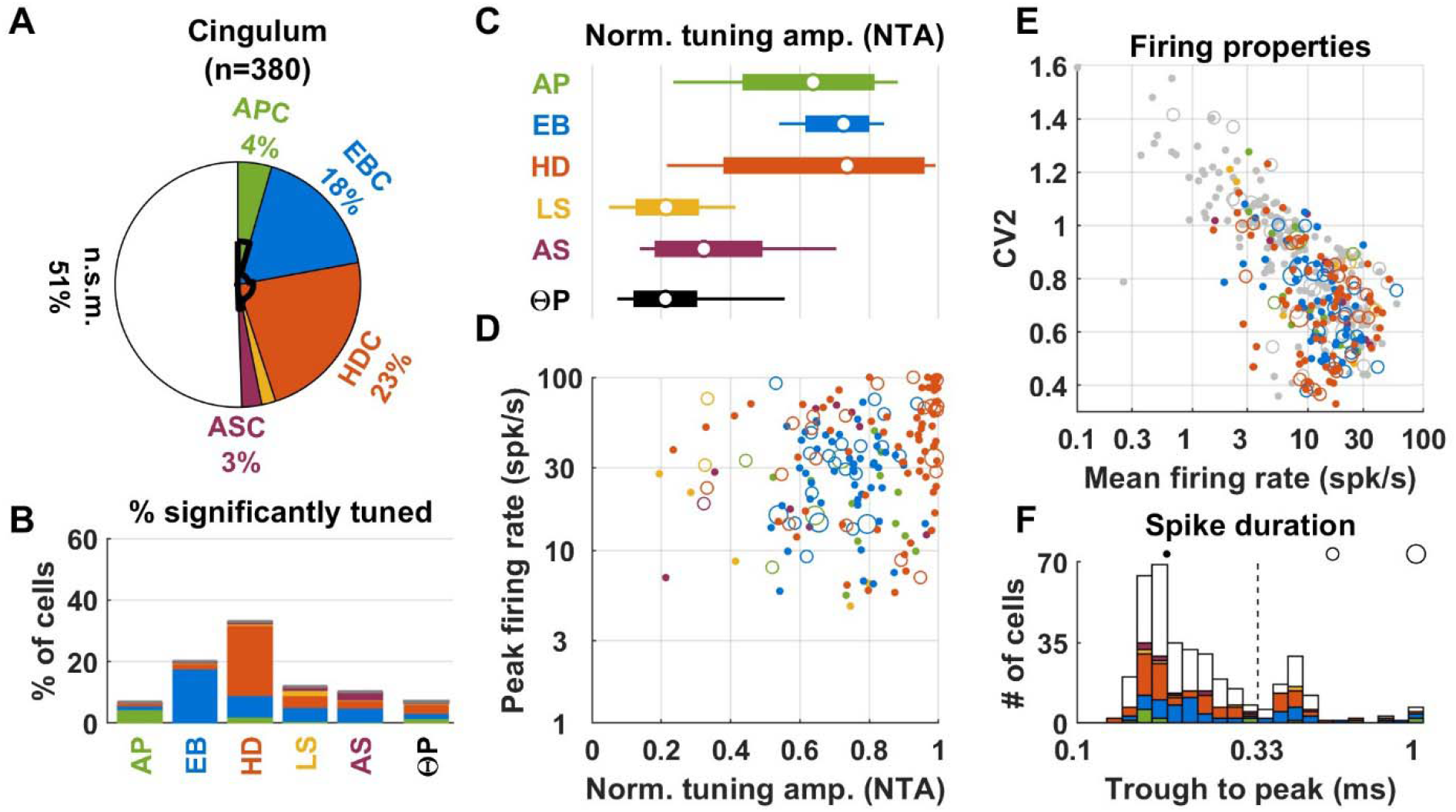
Population response in the cingulum. Same legend as in **Fig. 6**. Summary data from 5 of 7 animals (**Suppl. Fig. 9**), excluding H62M and H65M (show in **Suppl. Fig. 9**) because of minimal locomotor behavior. Cingulum responses resembled a mixture of ATN and RSC neurons, consistent with anatomical studies showing that it conveys anterior thalamic and RSC projections (Bubb et al. 2018). The majority of cells were classified as HDC (23%) or EBC (18%). For HD responses, the median NTA was 0.92 (C), similar (Wilcoxon rank sum test, p=0.16) to ATN (0.89), with overlapping range (0.57-0.99 1^st^-9^th^ decile). Because fewer neurons with low firing rate (e.g. ^~^3 spk/s or less) or low peak responses (e.g. ^~^10 spk/s or less) were encountered (compare panels D,E with **Fig. 6D,E**), mean firing of HDC was significantly higher in the cingulum (mean firing: median = 14 vs 9 spk/s, p=2.10^−3^, Wilcoxon rank sum test; peak firing: median = 48 vs 20 spk/s, p<10^−5^). The NRA of EBC cells ranged from 0.57 to 0.84 (1^st^-9^th^ decile, C), overlapping the distribution in RSC, with a similar median (0.75 vs. 0.7). Furthermore, 39% of cingulum EBC were also significantly tuned to HD, with median NRA=0.24 (B), in agreement with findings in RSC (**Fig. 7B**, p=0.4, Wilcoxon rank sum test). Cingulum EBC had larger firing rate (14 vs 8 spk/s, p<2.10^−3^) and peak firing (30 vs 16 spk/s, p<10^−4^) (compare panels D,E with **Fig. 7D,E**). Small fractions of APC, LSC and ASC were encountered in the cingulum (A,B), with similar properties as in other regions. However, only a marginal theta rhythm could be identified in the LFP (see also **Suppl. Fig. 9**). Accordingly, only a small fraction (7%) of neurons exhibited a significant ΘP modulation, and the amplitude of this modulation was very low (median: 0.21). The average firing rate and CV2 in the cingulum (E, Spearman rank correlation=-0.64, p<10^−10^) largely overlapped distributions in ATN and RSC. As expected, the majority of neurons (300/380, 79%) had short-duration spikes (F).

**Supplemental Figure 9:**
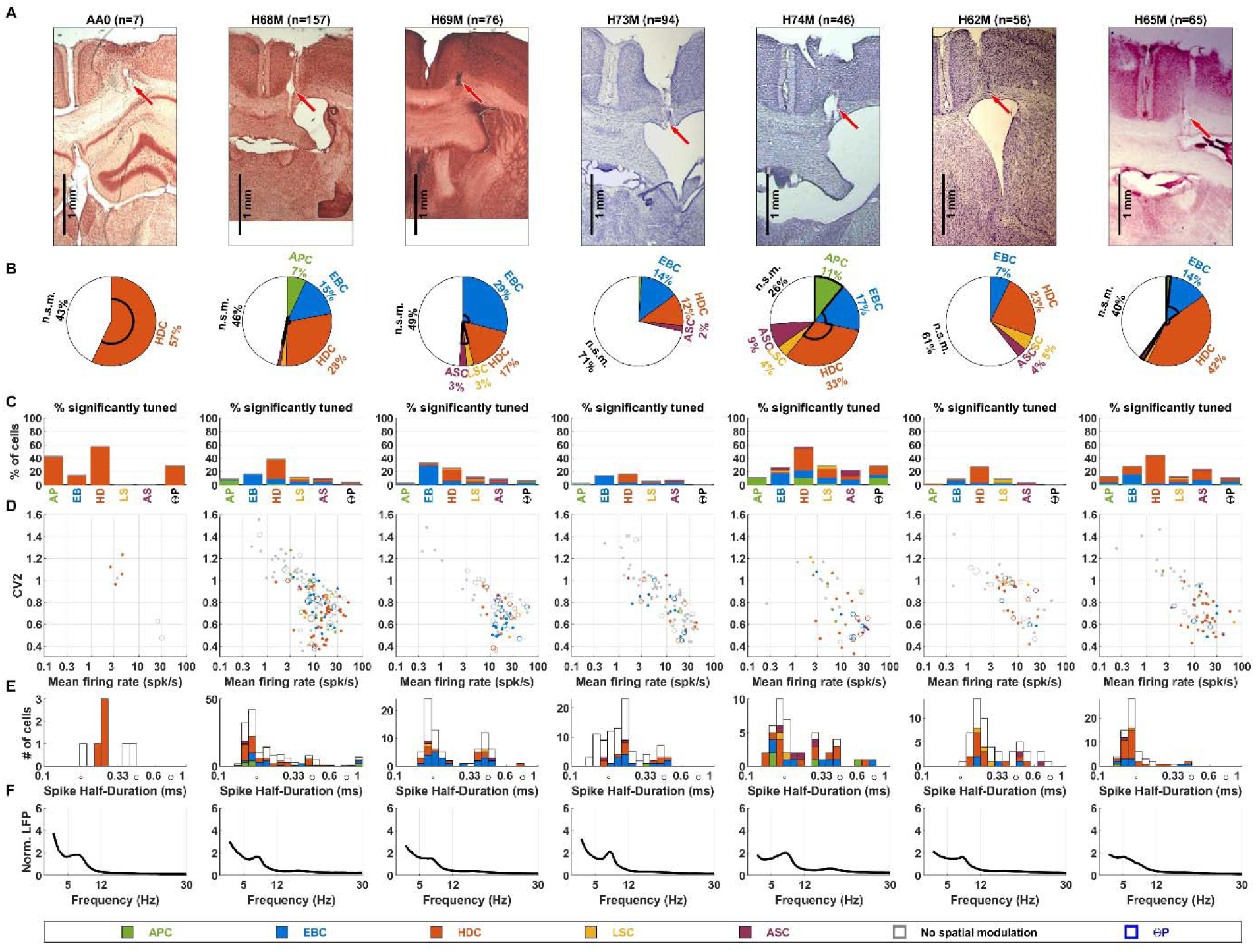
Recordings in the cingulum of individual mice. Same legend as in **Suppl. Fig. 2**. Animals H62M and H65M exhibited minimal locomotor behavior and were excluded from the population analysis since criteria for coverage of the arena interior were never met (but included here as LN analysis is robust to partial arena coverage).

**Supplemental Figure 10:**
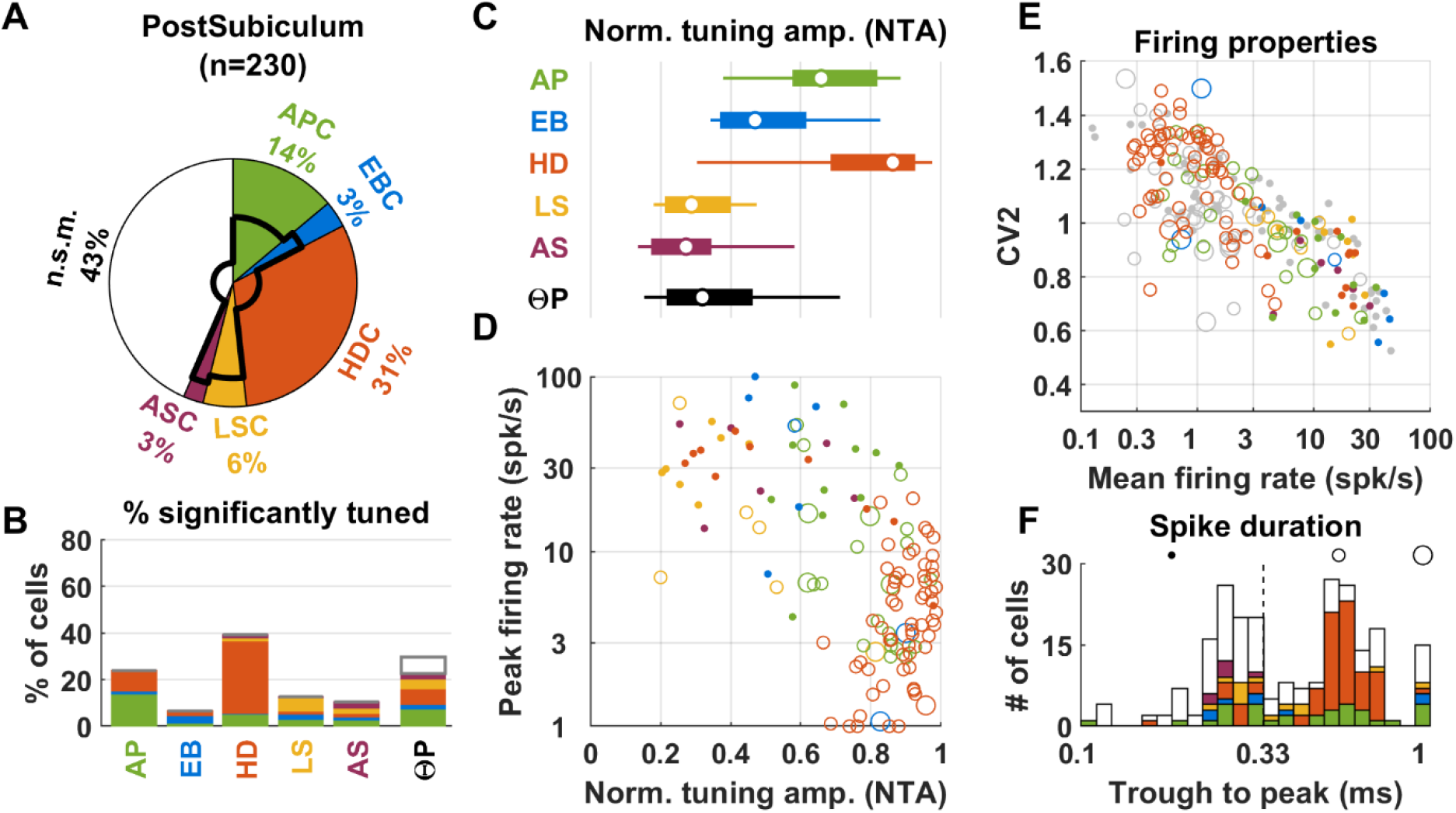
Population responses (3 animals, 230 neurons) in the Postsubiculum from previously published data (Peyrache et al. 2015). Same legend as in **Fig. 6**. The most prominent cell type (31%) was HDC (panel A,B). Most (61/71; 86%) HDC had long spike duration (panel F). These cells, which appear as large open orange symbols in panel D, had large NTA (median = 0.91; 1^st^-9^th^ decile: 0.77-0.97) and low peak firing (median = 3.8 spk/s; 1^st^-9^th^ decile: 1.2-10.5 spk/s), likely correspond to layer 3 pyramidal neurons identified as the main population of postsubicular HDC in (Tukker et al. 2015, Preston-Ferrer et al. 2016, Simonnet et al, 2017; Simonnet et Fricker, 2017). The second most prominent type (14%) was APC. Most (32/43, 74%) APC had long-duration spikes, and AP responses had generally large NTA (median = 0.67; 1^st^-9^th^ decile: 0.38-0.89). We noted that 12 APC were significantly tuned to HD and 20 HDC were significantly tuned to AP, such that, altogether, ^~^14% of the population encoded AP and HD conjunctively. Cells that encode these variables conjunctively were also reported by Caccuci et al. (2004), although most of these cells were Θ-modulated. In contrast, the conjunctive cells identified in our analysis of the Peyrache et al’s (2015) dataset were rarely (9/32, 28%) Θ-modulated.

**Supplemental Figure 11:**
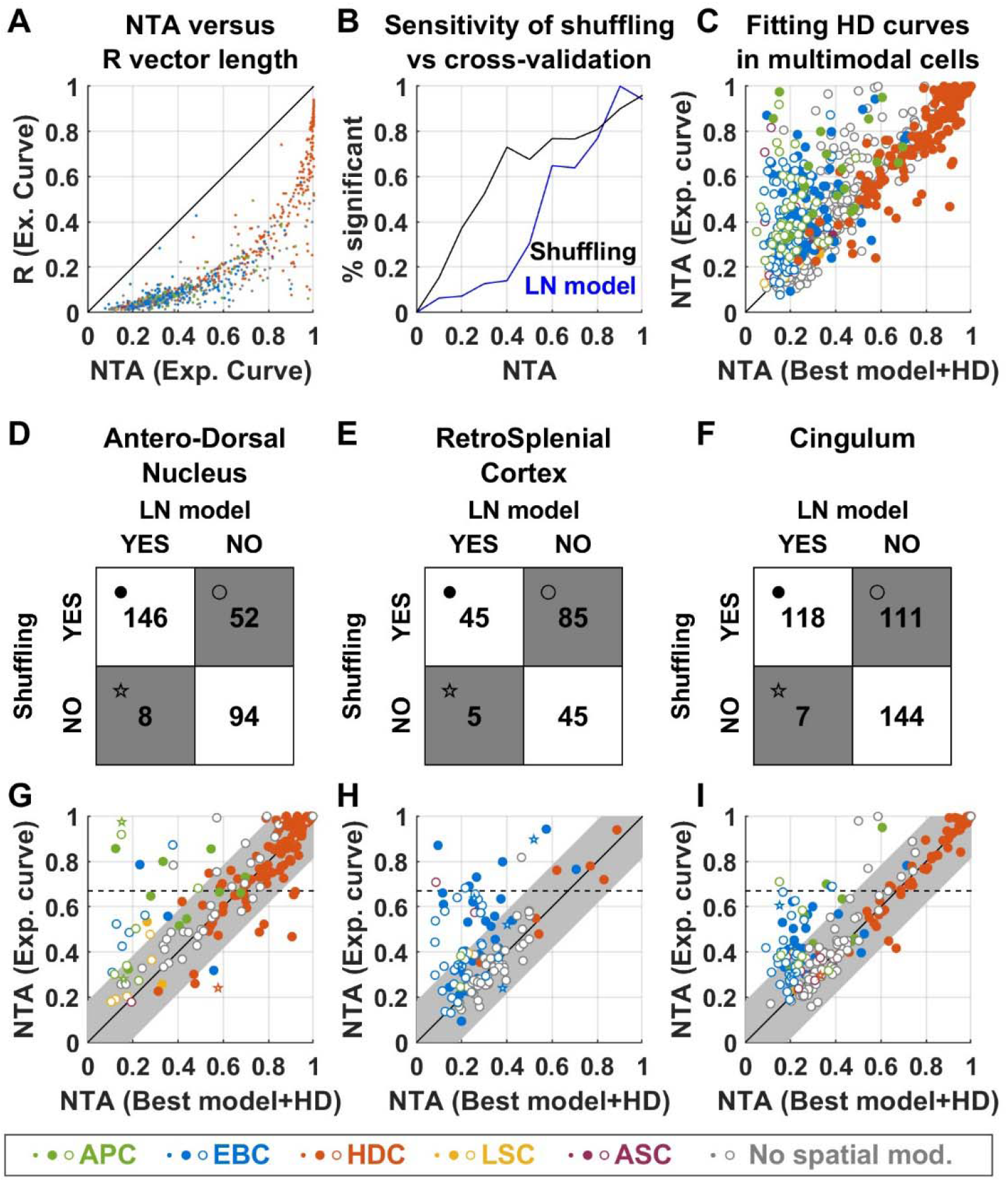
Classification of HD cells using the LN model versus traditional approaches. Here we analyze how the statistical approach used to classify HD cells in the LN model differs from traditional techniques where HD tuning is quantified by the mean vector length |R|, and statistical significance is evaluated by a shuffling test where |R| is considered significant if it is larger than 99% of a set of |R| values produced by randomly shifting the animal’s motion relative to the neuronal spike train (Skaggs 1993; Finkelstein et al. 2015). Alternatively, many studies classify cells as HDC when |R| exceeds a threshold, e.g. 0.26 (Jacobs et al. 2017) or 0.4 (Yoder and Taube 2009; Kornienko et al. 2018). In contrast, the LN model tests for HD tuning by fitting a HD tuning curve to 90% of the recorded data and measuring how the fitted curve accounts for the cell’s firing during the remaining 10%. This operation, called cross-validation, is repeated 10 times. A cell is considered HD tuned if the fitting quality is significantly larger than 0 (or than a previous model that doesn’t include HD) based on a signed rank test over these 10 samples. Another fundamental difference between the LN model and traditional approaches is that the LN model fits the cell’s firing with multiple variables simultaneously. Here, we discuss how these differences affect the classification of HD cells by considering data from the ATN, cingulum and RSC only (where most HD-tuned cells are found). **A:** Comparison between the traditional measure, |R|, and the NTA measure used in this study. We computed the |R| and NTA values of the experimental tuning curves of all cells (regardless of whether they were significantly tuned to HD). Most data points cluster tightly to form a curve, indicating that there is a close (although non-linear) correspondence between |R| and NTA. Cells are color-coded based on the classification by the LN model (see legend). **B:** Comparison between the sensitivities of the cross-validation and shuffling tests. We performed a shuffling test on the |R| value of all cells. Independently, we fitted a LN model where only HD was included. Next, we computed the percentage of cells classified as HD-tuned based on a shuffling test (black curve) or the LN model (blue curve) as a function of NTA. As expected, almost all cells pass both tests when the NTA is high (>0.8). In contrast, fewer cells with intermediate NTA (0.2-0.8 range) pass the LN model’s cross-validation test. Thus, the cross-validation procedure is less sensitive than the shuffling test. **C:** Previous studies (Muller et al. 1994; Cacucci et al. 2004; Rubin et al. 2014) have pointed out that responses to variables other than HD (e.g. AP or EB) can be erroneously interpreted as HD tuning. When this happens, HD tuning will appear high when fitting the LN model with only the HD variable, or when computing the experimental tuning curve, but will be lower when the response to other modalities is accounted for, as done by the LN model in general. To appreciate this, we plot the NTA of the experimental HD tuning curve versus the NTA of the HD curve fitted by the LN model. The latter was computed based on the cell’s best model (for HD-tuned cells) or by adding the HD variable to the best model. Filled/open symbols represent HD tuned/not tuned based on the full LN model, color-coded based on the classification by the LN model. Many AP and EB cells (green and blue) appear above the diagonal, indicating that responses identified as AP or EB are erroneously interpreted as HD tuning when considering only the HD model. This overestimation may happen in cells that are really HD tuned (closed symbols) or not (open symbols). From B and C, we conclude that (1) the shuffling test is more sensitive than the cross-validation procedure, and that (2) computing NTA (or equivalently |R|) from experimental tuning curves is prone to overestimating HD tuning in multimodal cells. Next, we examine how these approaches differ, in practice, when applied to ATN, RSC and cingulum. **D-F:** Contingency matrices indicating the number of cells classified as HD-tuned or not by the LN model and shuffling method in ATN (D), RSC (E) and cingulum (F). The symbols shown in the matrix correspond to the symbol code in panels G-I. The two methods generally agree in the ATN: 146 cells are classified as HD-tuned and 94 as HD non-tuned by both, i.e. the classification matches in 240/300 (80%) cells. 52 cells (17% of ATN) are classified as HD-tuned by only the shuffling method, and a negligible fraction by only the LN model. The two classifications diverge to a larger extent in RSC and cingulum, where 48% and 27% of cells are classified as HD-tuned based on the shuffling methods only. **G-I:** To elucidate the origin of these discrepancies, we plot the NTA of the experimental HD tuning curve versus the curve fitted by the LN model (as in panel C). Cells that are HD-tuned based on both classifications are shown as filled symbols. Cells that are HD-tuned based on the shuffling/cross-validation methods only are shown as open disks/stars. Cells are color-coded based on the classification by the LN model (see legend). A sizeable fraction of cells are classified as HD tuned based on the shuffling method only (open symbols, n=248 across all 3 areas), and we reason that this may occur for two reasons: (1) because the NTA of some cells is overestimated when computing the experimental curve (see panel C), in which case cells will appear above the diagonal and/or (2) because the shuffling test is more sensitive for cells with low NTA (see panel B). To quantify approximatively how many cells fall in each category, we estimate a confidence interval around the diagonal by considering cells that are classified as HDC by both methods (solid red). The confidence interval width is set at the 95% percentile of the distance distribution of these cells from the diagonal (0.13). We find that 193/248 (78%) of open symbols fall within this interval. These likely correspond to cells where HD tuning was likely *not* over-estimated, and that were classified as HD-tuned because of the shuffling test’s larger sensitivity. These cells are likely genuinely HD-tuned, although with a low amplitude. In contrast, HD tuning may have been overestimated in cells that appear above the interval (55/248; 22% of open symbols), and the classification as HD-tuned may be erroneous. In total, this category of potentially mis-classified cells represents 16/300 (5%) ATN cells, 18/180 (10%) RSC cells and 19/380 (5%) cingulum cells. Finally, we note that some studies require HD cells to pass a threshold, i.e. |R|>=0.26 in (Jacobs et al. 2017) or |R| >=0.4 (Yoder et al. 2009; Kornienko et al. 2018). Based on panel A, we estimate that |R|>0.26 corresponds to NTA>=0.67 (broken lines in G-I). When this threshold is added to the shuffling test, a total of 223 cells are classified as HD-tuned, out of which 192 (86%) are also classified as HD-tuned by the LN model. However, this test now rules out 137 out of 329 cells (42%) that are classified as HD-tuned by the LN model, including, in particular, most (77/91, 85%) cells identified as APC or EBC with significant HD tuning by the LN model. Thus, using a threshold allows selecting well-tuned HD cells in a conservative manner, but tends to miss weaker HD cells and multimodal cells.

**Supplemental Figure 12:**
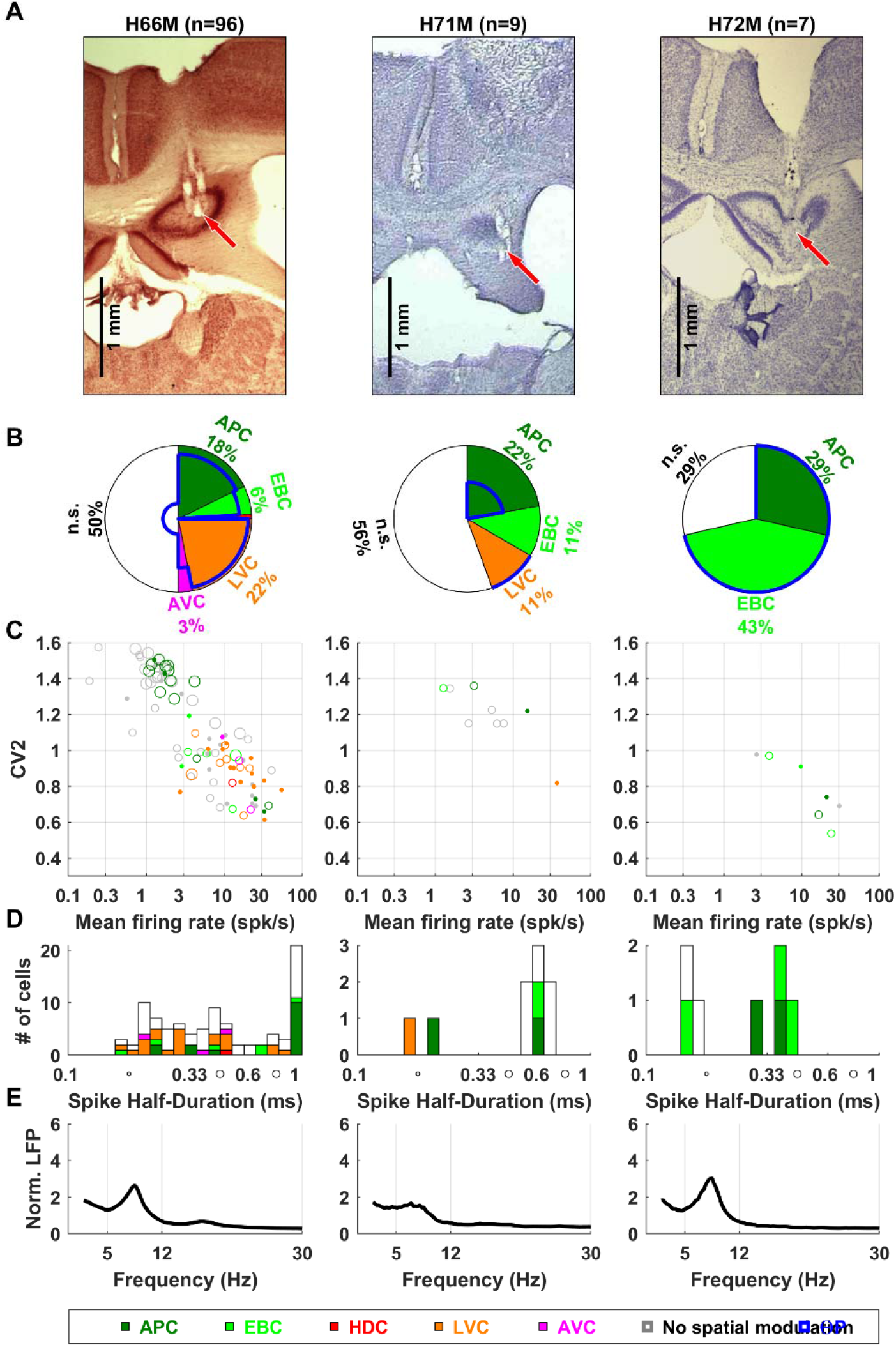
Recordings in the hippocampus of individual mice. Same legend as in **Suppl. Fig. 2**.

**Supplemental Figure 13:**
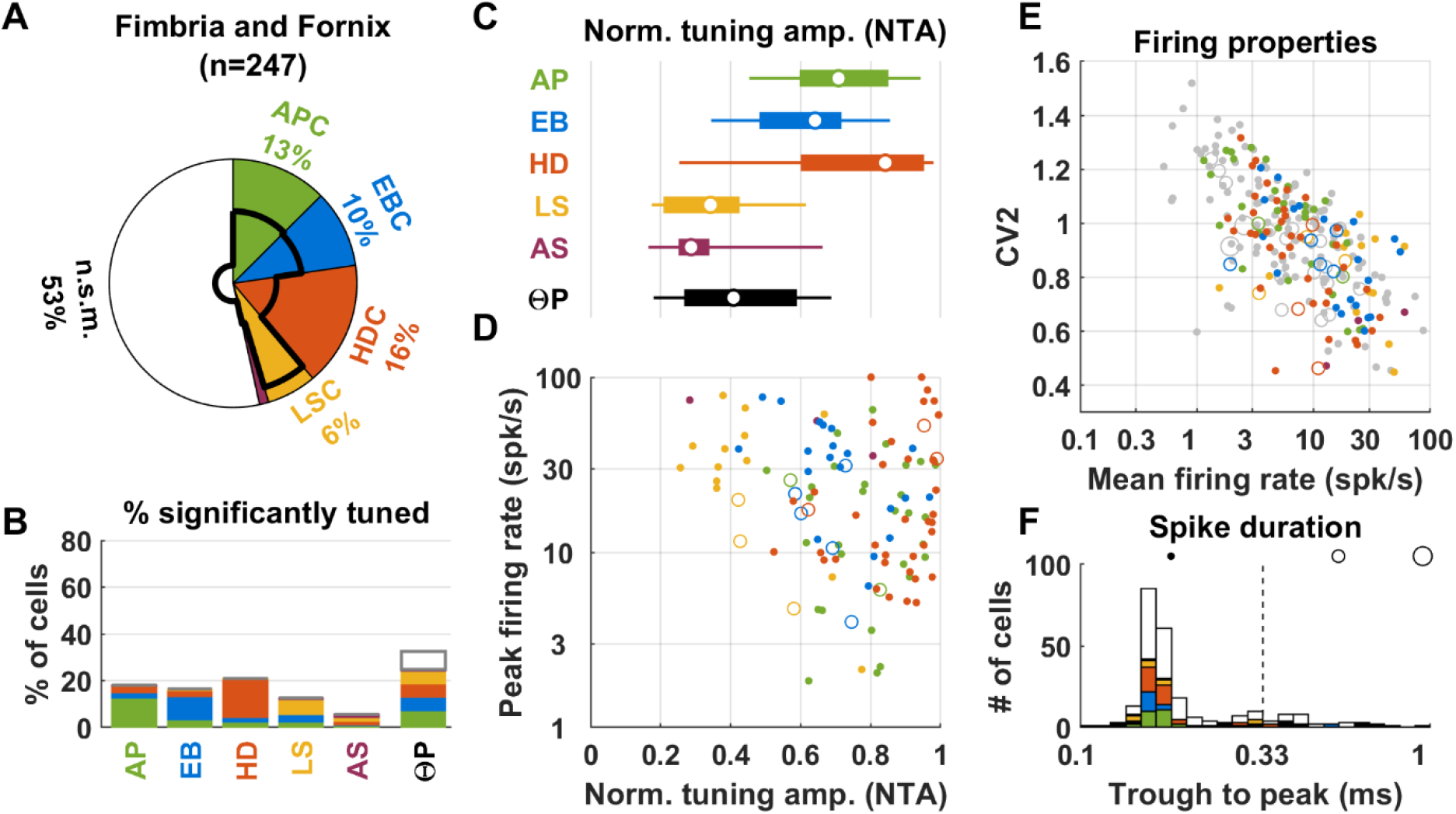
Population responses in the fimbria and fornix. Same legend as in **Fig. 6**. A summary of population responses follows. *APC:* A fraction (13%) of cells recorded in the fimbria and fornix were classified as APC (panels A,B). APC in the fimbria exhibited large NTA (panels C,D, median = 0.81, 1^st^-9^th^ decile: 0.62-0.95); as well as large peak firing rate (median 16 spk/s, 1^st^-9^th^ decile 3-36 spk/s; D). *HDC:* A fraction (16%) of cells recorded in the fimbria and fornix were HDC (panels A,B). The median NTA was high (panels C,D; median: 0.91, 1^st^-9^th^ decile 0.65-0.98) and similar as in ATN and cingulum (p=0.49 and p=0.62 respectively). The median peak firing rate (16 spk/s; 1^st^-9^th^ decile: 6-73) was similar to the value observed in the ATN (p=0.33, Wilcoxon rank sum test) and cingulum (p=0.7). *Other cells: We* also encountered a sizeable fraction of EBC (10%, panel A) that had large NTA (median 0.69, range 0.54-0.9), similar to RSC (p=0.17); as well as a fraction of LSC (6%) with moderate NTA (median 0.42, 1^st^-9^th^ decile: 0.3-0.69) similar to hippocampal LSC (p=0.12). *Theta rhythm:* Across the entire population of recorded neurons, 32% of fimbria neurons were ΘP modulated. However, this fraction increased to 53% when only spatially modulated neurons were considered. *Spiking properties:* Mean firing rate and CV2 were inversely correlated, as in other regions (panel E, Spearman rank correlation=-0.65, p<10^−10^). Most cells (86%) had short duration spikes, as expected from recordings performed in the white matter.

**Supplemental Figure 14:**
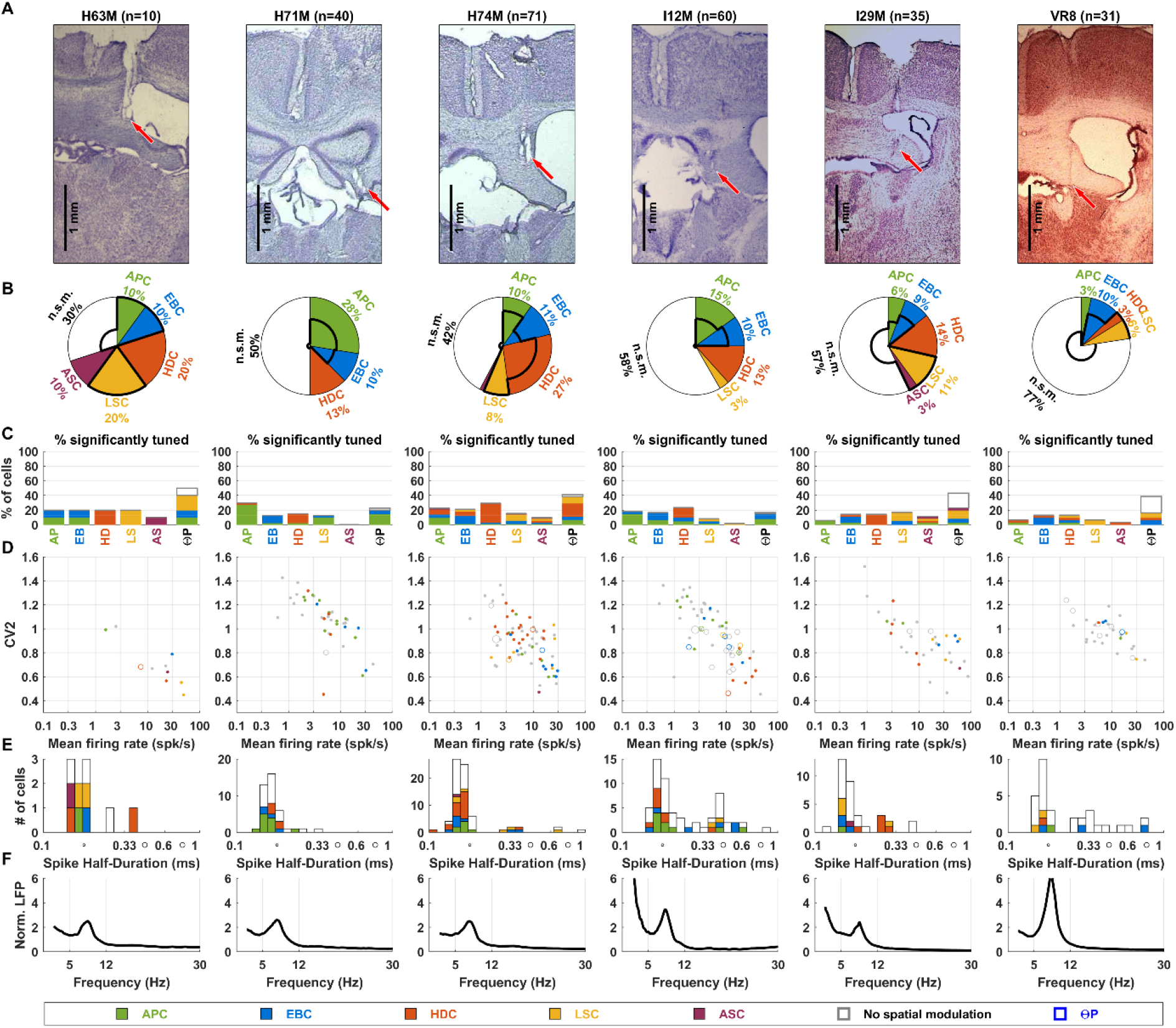
Recordings in the fimbria and fornix of individual mice. Same legend as in **Suppl. Fig. 2**.

**Supplemental Figure 15:**
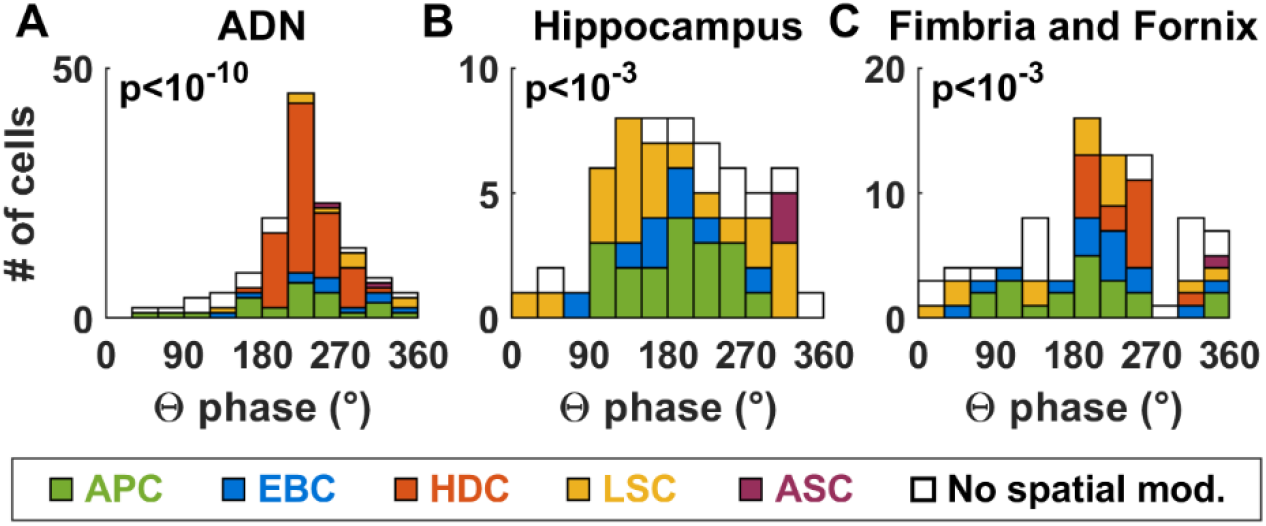
Distribution of preferred phase of ΘP-modulated cells. Distributions are shown as histograms, where phases of 0, 90, 180 and 270° correspond to the trough, ascending phase, crest and descending phase of the LFP. Cells are color-coded based on their classification. About half of ATN and hippocampal cells, and half of spatially-modulated neurons recorded in the fimbria, responded preferentially at a certain phase of the Θ-band LFP. We found that the distributions of preferred phase were non-uniform. In ATN (**A**), most cells responded preferentially during the descending phase (180-360°), with an average preferred phase of 230±10°. In particular, the two main classes of ΘP-modulated ATN neurons, HDC and APC, had an identical average preferred phase (230 vs 235°, p=0.7, Watson-Williams test). In the hippocampus (**B**), the average preferred phase was 193±29° amongst APC, whereas preferred phases were distributed uniformly amongst LSC (yellow). In the fimbria (**C**), HDC responded preferentially in the descending phase (238±20°) whereas APC fired closer to the LFP crest 184±40° (p=5.10^−4^ versus ATN, Watson-Williams test).

**Supplemental Figure 16:**
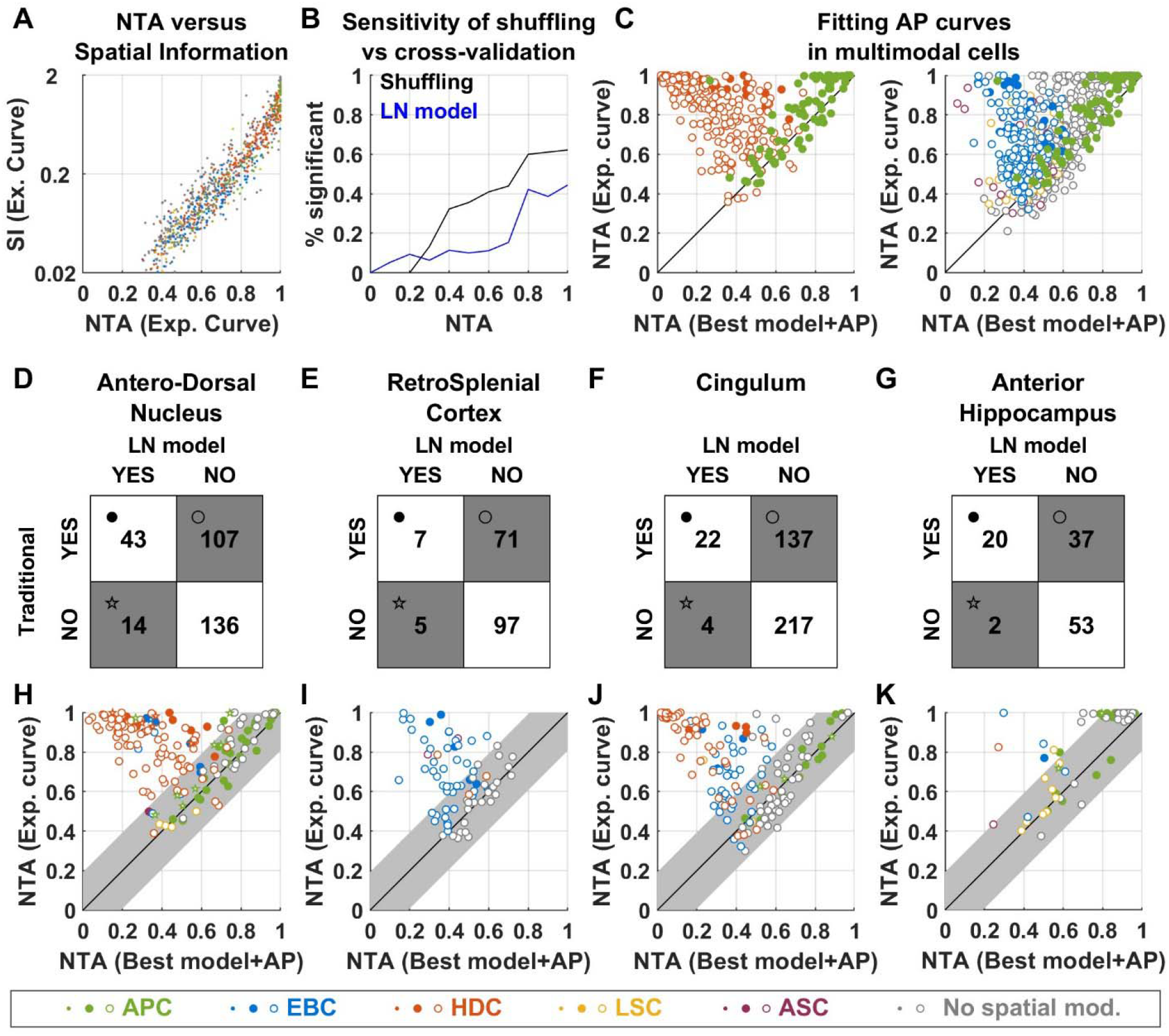
Classification of AP cells using the LN model versus traditional approaches. We now follow the same logic as in **Suppl. Fig. 13** to analyze how the statistical approach used to classify AP cells in the LN model differs from traditional techniques, where AP tuning is quantified by spatial information (SI) of the tuning curve, and where a give SI value is considered significant if it is larger than 99% of a set of shuffled values (Skaggs 1993; Rubin et al. 2014). We consider data from the ATN, cingulum and anterior hippocampus only (where most AP-tuned cells are found). **A:** Comparison between spatial information (SI), and the NTA measure used in this study. We plot the SI and NTA of the experimental tuning curves of all cells (regardless of whether they were significantly tuned to AP). Most data points cluster tightly to form a curve, indicating that there is a close correspondence between SI and NTA. Cells are color-coded based on the classification by the LN model (see legend). **B:** Comparison between the sensitivities of the cross-validation and shuffling tests, as in **Suppl. Fig. 13B**. Even when NTA is high (>0.8), only ^~^60% and ^~^40% cells pass the shuffling or cross-validation test, respectively. This indicates that apparently high AP tuning may occur randomly; but that these occurrences will be classified as non-significant based on both the shuffling test or LN model. This didn’t occur with HD tuning (**Suppl. Fig. 13B**): this difference may be due to the fact that repetitively covering the 2D surface of the arena, which is required to archive statistical robustness, is harder than covering the 1D space of head direction. Similar to HD tuning, we find that the cross-validation procedure is generally less sensitive than the shuffling test for intermediate NTA. **C:** In multimodal cells, responses to variables other than AP (e.g. HD) can be erroneously interpreted as AP tuning using the traditional analysis. We plot the NTA of the experimental AP curve versus the NTA of the AP curve fitted by the LN model. For readability, we separate this panel in two plots, with HD cells and AP cells on the left and AP cells and other cell types on the left. Filled/open symbols represent cells that are HD tuned/not tuned based on the full LN model. Cells are color-coded based on the classification by the LN model (see legend). We find a striking number of HD cells (red) converging towards the upper left corner, i.e. where AP tuning appears very strong if HD tuning is not accounted for first. Thus, as pointed out by previous studies (Peyrache et al. 2017), HD cells may easily be confounded for AP cells. We also find that AP tuning is often overestimated in EB cells (blue). **D-G:** Contingency matrices indicating the number of cells classified as AP-tuned or not by the LN model and shuffling method in ATN (D), RSC (E), cingulum (F) and anterior hippocampus (G). We first note that most cells that were classified as AP-tuned by the LN model are also classified as AP-tuned by the shuffling test (ATN: 43/57, i.e. 75%; cingulum: 22/26, i.e. 85%; hippocampus: 20/22, i.e. 91%). This validates our finding that the ATN and cingulum contain APC populations. Next, we observe that the ATN, RSC and cingulum contain large fractions (36%, 39% and 36% respectively) of cells that are incorrectly classified as AP-tuned by the shuffling test only (open circles). **H-K:** As in **Suppl. Fig. 13G-I**, we plot the NTA of the experimental AP curve versus the curve fitted by the LN model. Cells tuned based on both classifications are shown as filled symbols. Cells tuned based on the shuffling/cross-validation methods only are shown as open disks/stars. Cells are color-coded based on the classification by the LN model (see legend). We draw a confidence interval, with a width estimated at 0.14 (based on cells classified as APC by both methods) around the diagonal. In the ATN (panel H), 68 cells are incorrectly classified as AP-tuned based on the shuffling procedure only and positioned above the confidence interval. Most of these cells (63/68) are HD cells (open orange circles). In the RSC (panel I), we also find a large group cells above the confidence interval, most of which (26/29) are EBC (open blue symbols). Thus, 32% (26/81) of RSC EBC would be erroneously characterized as APC. We also find a sizeable group of open symbols within the confidence interval (42 cells, i.e. 59% of cells classified as AP based on the shuffling method only). This suggests that a population of cells with weak but significant AP tuning, that would not have been detected by the LN model, may exist in the RSC. In the cingulum (panel J), we find 45% of HDC and 40% of EBC above the diagonal; as well as a group of cells within the confidence interval (47% of cells classified as AP based on the shuffling method only). These results are in line with the hypothesis that cingulum carries a mixture of ATN and RSC signals. In total, 68/300 (23%) ATN cells, 29/137 (16%) RSC cells and 72/380 (19%) cingulum cells, placed above the diagonal, were incorrectly classified as AP-tuned by the shuffling test. In the hippocampus (panel K), most (30/37) open symbols falls within the confidence interval. Furthermore, many cells cluster at the upper right corner of the graph, indicating that they are highly tuned to AP even when this tuning is overestimated by the experimental curve. This indicates that most cells identified by the shuffling test may be genuine AP cells, and that the LN model may have lacked the sensitivity to identify them. A possible remedy for this lack of sensitivity could be to perform longer recording sessions.

**Supplemental Movie 1: Response of a RSC EBC (same as in Fig. 5) during free exploration**. The movie shows the animal’s motion and neuronal activity recorded during two minutes of free exploration. **Upper left panel:** animal motion from an allocentric point of view. The arena’s boundary is shown in white, with the cue card represented as a dark gray arc. As time elapses, the animal’s trajectory (light blue) and neuronal spikes (red dots) are shown. The arrow indicates the position of the nearest wall. The **right panel** displays exactly the same image, except for the light blue trajectory and red dots. The image is rotated in order to appear in egocentric coordinates. The light blue trajectory and red dots now represent the trajectory of the nearest boundary in egocentric coordinates. Spikes concentrate in front of the head, indicating that the neuron responds when the head is close to the wall and faces it. Lower left panel: raw neuronal data.

**Supplemental Data 1:** Classification and response of all cells included in this study. Data is organized as a spreadsheet. For each cell, we provide the NTA and peak response to all variables (values are set to −1 for variables that don’t modulate the cell significantly), as well as the cell’s mean firing rate, CV2 and trough to peak spike duration.

